# Distinct Heterogeneity in the Naive T cell Compartments of Children and Adults

**DOI:** 10.1101/2022.10.04.510869

**Authors:** Claire E. Gustafson, Zachary Thomson, Ziyuan He, Elliott Swanson, Katherine Henderson, Mark-Phillip Pebworth, Lauren Y. Okada, Alexander T. Heubeck, Charles R. Roll, Veronica Hernandez, Morgan Weiss, Palak C. Genge, Julian Reading, Josephine R. Giles, Sasikanth Manne, Jeanette Dougherty, CJ Jasen, Allison R. Greenplate, Lynne A. Becker, Lucas T. Graybuck, Suhas V. Vasaikar, Gregory L. Szeto, Adam K. Savage, Cate Speake, Jane H. Buckner, Xiao-jun Li, Troy R. Torgerson, E. John Wherry, Thomas F. Bumol, Laura A. Vella, Sarah E. Henrickson, Peter J. Skene

## Abstract

The naive T cell compartment undergoes multiple changes across age that associate with altered susceptibility to infection and autoimmunity. In addition to the acquisition of naive-like memory T cell subsets, mouse studies describe substantial molecular reprogramming of the naive compartment in adults compared with adolescents. However, these alterations are not well delineated in human aging. Using a new trimodal single cell technology (TEA-seq), we discovered that the composition and transcriptional and epigenetic programming of the naive T cell compartment in children (11-13 yrs) is distinct from that of older adults (55-65 yrs). Naive CD4 T cells, previously considered relatively resistant to aging, exhibited far more pronounced molecular reprogramming than naive CD8 T cells, in which alterations are preferentially driven by shifts in naive-like memory subsets. These data reveal the complex nature of the naive T cell compartment that may contribute to differential immune responses across the spectrum of human age.

**One Sentence Summary:** The naive CD8 and CD4 T cell compartments in humans are heterogeneous and impacted differently with age, in which naive CD8 T cell subsets dramatically shift in composition and true naive CD4 T cells display significant molecular re-programming.

## Main Text

### INTRODUCTION

At the extremes of age, there are alterations in immune responses that lead to increased morbidity and mortality from many infections, such as Influenza A and *Streptococcus pneumoniae*. However, these alterations are not identical in children and adults, and can be observed in the context of age-specific susceptibility to certain pathogens. This phenomenon is particularly highlighted in the on-going global pandemic of SARS-CoV-2, in which older adults have dramatically higher rates of hospitalizations and death from infection than children. (*1*) Naive T cell responses are critical for developing defenses against emerging viral infections and long-lasting, effective vaccine responses. However, the changes in this compartment that contribute to differential immunity between healthy children and adults are not well defined.

A consistent hallmark of T cell aging is the loss of naive CD8 T cells. Studies have also demonstrated that the naive CD8 T cell compartment is impacted by naive-like memory cell infiltration (*2*–*4*) as well as pseudo-differentiation of naive cells towards more memory-like epigenetic programming. (*5*) Mouse aging studies demonstrate that naive CD8 T cells have distinct developmental pathways, altered epigenetic programming and unique responses to infection in infancy compared with adulthood. (*6*, *7*) Conversely, the naive CD4 T cell compartment appears much less affected by age, showing less decline in numbers and fewer molecular alterations across adult aging. (*8*) However, naive CD4 T cells still displays functional differences in antigen-specific response in children and adults, with children preferentially polarizing towards T_H_2 programming compared with adults. (*9*, *10*) Differential polarization suggests that naive CD4 T cells in children and adults may have distinct molecular programming at a resting state, similar to that observed in mouse naive CD8 T cell populations. However, the contribution of compositional versus intracellular reprogramming across human age is unclear. A detailed analysis of cellular and molecular heterogeneity of human naive CD8 and CD4 T cells is needed to better understand the differential immune responsiveness observed in children and older adults.

Using single cell sequencing techniques, it has been demonstrated that the composition of the mouse T cell compartment significantly changes with age. (*6*, *7*, *11*, *12*) These comprehensive studies are still limited in humans. Moreover, most studies have been restricted to analysis of cellular heterogeneity using a single modality at a time (i.e., RNA or ATAC or protein), with limited ability to deconvolute complex cellular alterations. A newly developed trimodal single cell assay (TEA-seq) permits the simultaneous single cell analysis of surface protein, RNA and chromatin accessibility. (*13*) This trimodal approach is of particular importance because certain canonical markers can only be assessed in one modality and are often lost in single-modality analyses, particularly when looking at resting, non-activated T cells, such as gene isoforms (e.g., CD45RA by antibody), cytokines (which often require cell stimulation in more naive-like populations) and transcription factor expression (which requires complex intracellular staining for protein detection). The ability to differentiate T cell subsets via these three modalities together provides a unique opportunity to directly study the interplay between canonical surface protein phenotypes and the transcriptional and epigenetic programs within cells, thus allowing for unprecedented and detailed resolution of the complex heterogeneity within the naive T cell compartment in humans. In this study, we utilize TEA-seq to dissect the impact of compositional and molecular alterations within the naive T cell compartment in children and adults. From these analyses, we identify key changes in naive T cell heterogeneity that may underlie differential immune capacity across age. In support of open science and to enable the research community to discover additional novel insights from this rich dataset, we also provide a user-friendly data visualization tool at: https://explore.allenimmunology.org/explore/db8f8010-674e-4e91-a0cb-a7b3123f9041.

## RESULTS

### Age significantly impacts the molecular landscape of phenotypically naive T cells

To enhance our understanding of naive T cell biology across the human age spectrum, we performed a deep multi-omic analysis of T cells isolated from the blood of pediatric (11-13 yrs, n=8) and adult (55-65 yrs, n=8) female donors using our recently developed tri-modal single cell assay TEA-seq. (**Figure 1A**) We analyzed a total of 300,418 single-positive T cells across all donors, composed of 204,586 CD4+ T cells and 95,832 CD8+ T cells. The historical gold standard for T cell subset characterization is flow cytometry using hierarchical gating based on a select set of surface markers and subsequent cell sorting. Therefore, we used a surface protein-based approach, as detected by oligo-tagged antibodies (antibody derived tags, ADTs), to perform cell gating analogous to flow cytometry. T cell subsets were defined by the expression of seven markers: CD4, CD8, CD127, CD25, CD45RA, CD27 and CCR7. (**Figure 1B, Supp Table 1**) Using the same gating approach as our flow cytometry dataset (**Supp Fig 1**), we determined that the frequencies of ADT-defined T cell subsets in our TEA-seq dataset were highly correlated with those of flow cytometry-based frequencies across all donors. (**Supp Fig 2A**) Changes in cell type frequency were consistent with the previously reported loss of peripheral naive CD8 T cells as one of the hallmarks of immune aging. (*14*) ADT-defined (CD45RA+CCR7+CD27+) naive CD8 T cells were significantly decreased in adults compared with pediatric donors (q=0.001, 26.1% pediatric vs 7.3% adult, mean of total T cells) (**Supp Fig 2B**). No significant decrease in naive CD4 T cells was observed. (q=0.29, 37.6% pediatric vs 30.1% adult, mean of total T cells). T cell subsets defined by ADTs differed substantially from those identified by RNA-based or ATAC-based label transfer methods based on comparison to reference datasets, with an average deviation of 29.3%. (**Supp Fig 3**) Thus, ADT-based cell labeling in TEA-seq is most reflective of the T cell populations defined by canonical flow cytometry analyses

**Figure 1.**
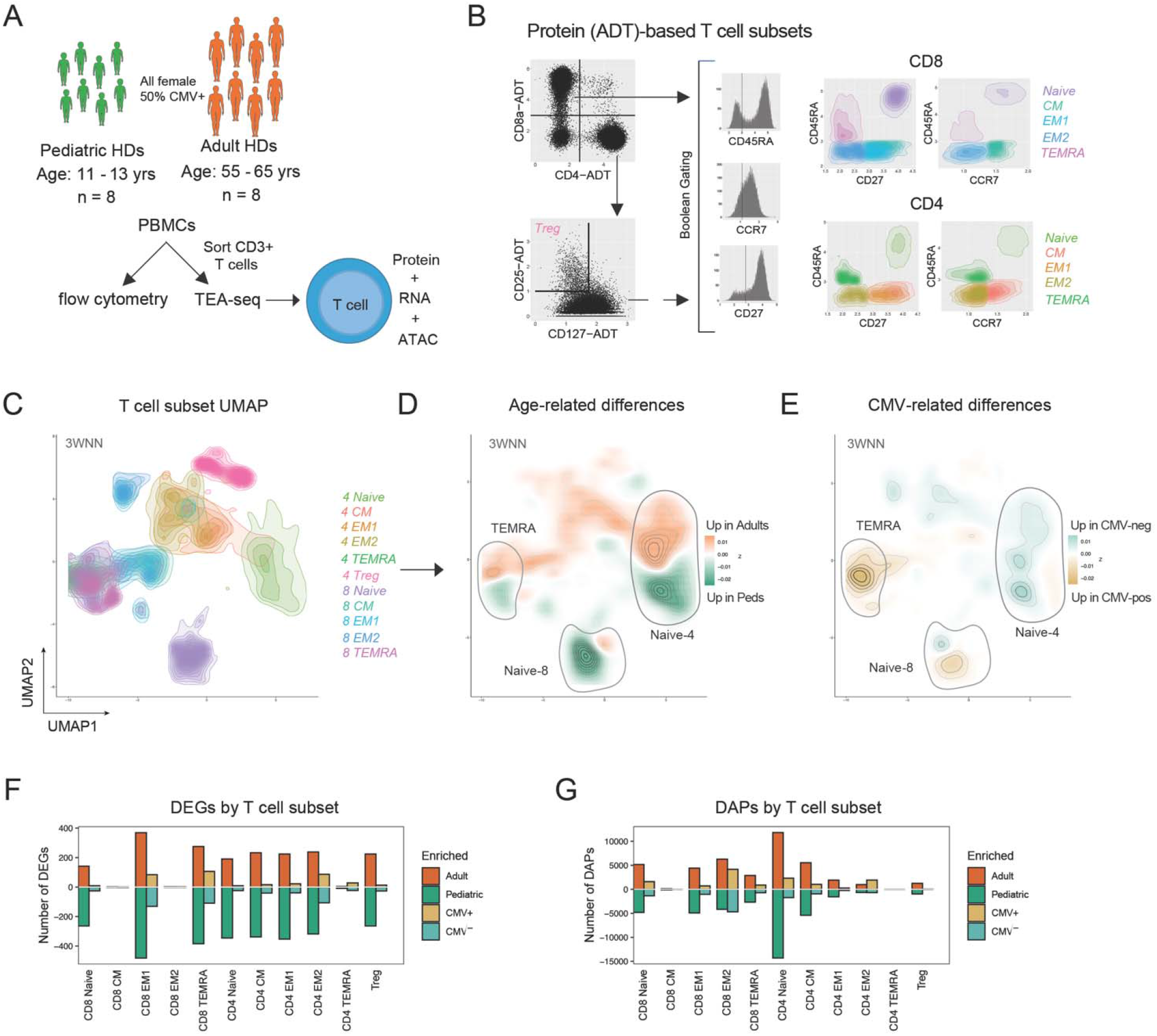
Age impacts the transcriptional and epigenetic landscape of T cell subsets. **(A)** Cohort and experimental overview. HDs, healthy donors. **(B)** T cell subset gating strategy using ADT expression. **(C)** 3WNN UMAP of ADT-defined T cell subsets from all donors, based on cell density. **(D)** UMAP showing change in cell density by age (green = greater in pediatric; orange = greater in adult) or **(E)** by CMV infection status (blue = greater in CMV-negative; yellow = greater in CMV-positive). **(F)** Number of differentially expressed genes (DEGs) and **(G)** number of differentially accessible peaks (DAPs) within each ADT-based T cell subset by age (green = up in pediatric; orange = up in adult) or CMV (blue = up in CMV-negative; yellow = up in CMV-positive).

In adults, aging can drive significant transcriptional changes within the T cell compartment (*14*). Thus, we first compared global transcriptional and epigenetic features across naive T cells in children and adults. For visualization, we generated a joint 2-dimensional UMAP projection of the multimodal data using a 3-way (ADT, RNA and ATAC combined) weighted nearest neighbor (3WNN) method. (*15*) T cell subsets are highlighted in **Figure 1C**. We observed age-specific changes, in which regions of the UMAP with higher cell density from adults are in orange and regions with higher cell density from pediatric donors are in green. Age corresponded with differences in cell distribution within cell types (e.g., within naive T cells) rather than frequency shifts across subsets (e.g., from naive to memory). (**Figure 1D**) For example, we see the naive CD4 T cell subset formed two distinct groups within the total naive CD4 T cell area, with one area of increased pediatric cell density and one of increased adult cell density. This high-level visualization suggests that there are both compositional changes between subsets and molecular changes within subsets with age.

The prevalence of CMV infection correlates with age and significantly impact the T cell compartment. (*16*, *17*) To enable us to compare age-related changes with those induced by CMV infection, our cohort was equally balanced by age and CMV infection status. Large shifts in cell type composition were correlated with CMV infection status, as highlighted by increased proportion of effector populations. (**Figure 1E**) CD8 TEMRA cells are predominantly from CMV-positive individuals, as indicated in yellow. This is consistent with elevated frequencies of both CD8 and CD4 TEMRA populations in CMV-positive individuals independent of age, in contrast to naive CD8 T cells that display a decrease in frequency with advancing age but not CMV infection status. (**Supp Fig 4**)

To further interrogate potential molecular changes in T cell subsets specific to age, we compared differential gene expression (using MAST) and chromatin accessibility (using ArchR) across each subset by age or CMV infection status. (**Figure 1F-1G**) Both at a transcriptional and epigenetic level, age had a greater impact on the number of differentially expressed genes (DEGs) and differentially accessible peaks (DAPs) than CMV infection status. Age induced elevated numbers of DEGs and DAPs changes across multiple subsets and differentiation states, including both naive CD8 and CD4 T cell subsets. CMV infection status had little impact on transcription and chromatin landscape in naive T cells, consistent with previous reports of CMV infection driving the expansion of effector memory populations but not naive or central memory populations. (*18*) These results demonstrate that age is associated with both transcriptional and epigenetic remodeling across the naive T cell compartment, whereas CMV infection status was associated with transcriptional alterations in effector populations with little epigenetic remodeling.

### Multiple memory-like subsets within phenotypically naive CD8 and CD4 T cell compartments

A subset of memory CD8 T cells can re-acquire a naive-like phenotype over the course of a lifetime. (*2*–*4*) We identified distinct sub-populations present within the “naive” CD8 and CD4 T cell compartments, using a combination of ADT, RNA and ATAC data in our TEA-seq dataset (**Figure 2A**) by performing unsupervised re-clustering of our ADT-defined naive (CD45RA^+^CCR7^+^CD27^+^) CD8 and CD4 T cells (45,481 and 99,434 total single cells analyzed, respectively) based on 3WNN. (**Figure 2B-2C**) Using the strategy outlined in **Figure 2A** for cell subset definitions, we identified 5 major cell subsets within the naive CD8 T cell compartment: true naive (CD49d-ADT^neg^CD95-RNA^neg^IFNG-ATAC^neg^), stem cell memory (SCM, CD49d-ADT^+^CD95-RNA^+^IFNG-ATAC^+^), 2 populations of memory-like naive precursors (MNP-1 and MNP-2, CD49d-ADT^high^CD95-RNA^low^IFNG-ATAC^low/+^), and mucosal-associated invariant T cells (MAIT, TCRVa7.2-ADT^+^ and CD161-ADT^+^). (**Figure 2D-2F**) We also identified 3 major cell subsets within the naive CD4 compartment: true naive, SCM and CD25^neg^ Tregs (FOXP3-RNA^+^CD25-ADT^neg^IL2RA-RNA^+^). (**Figure 2G-2I**)

**Figure 2.**
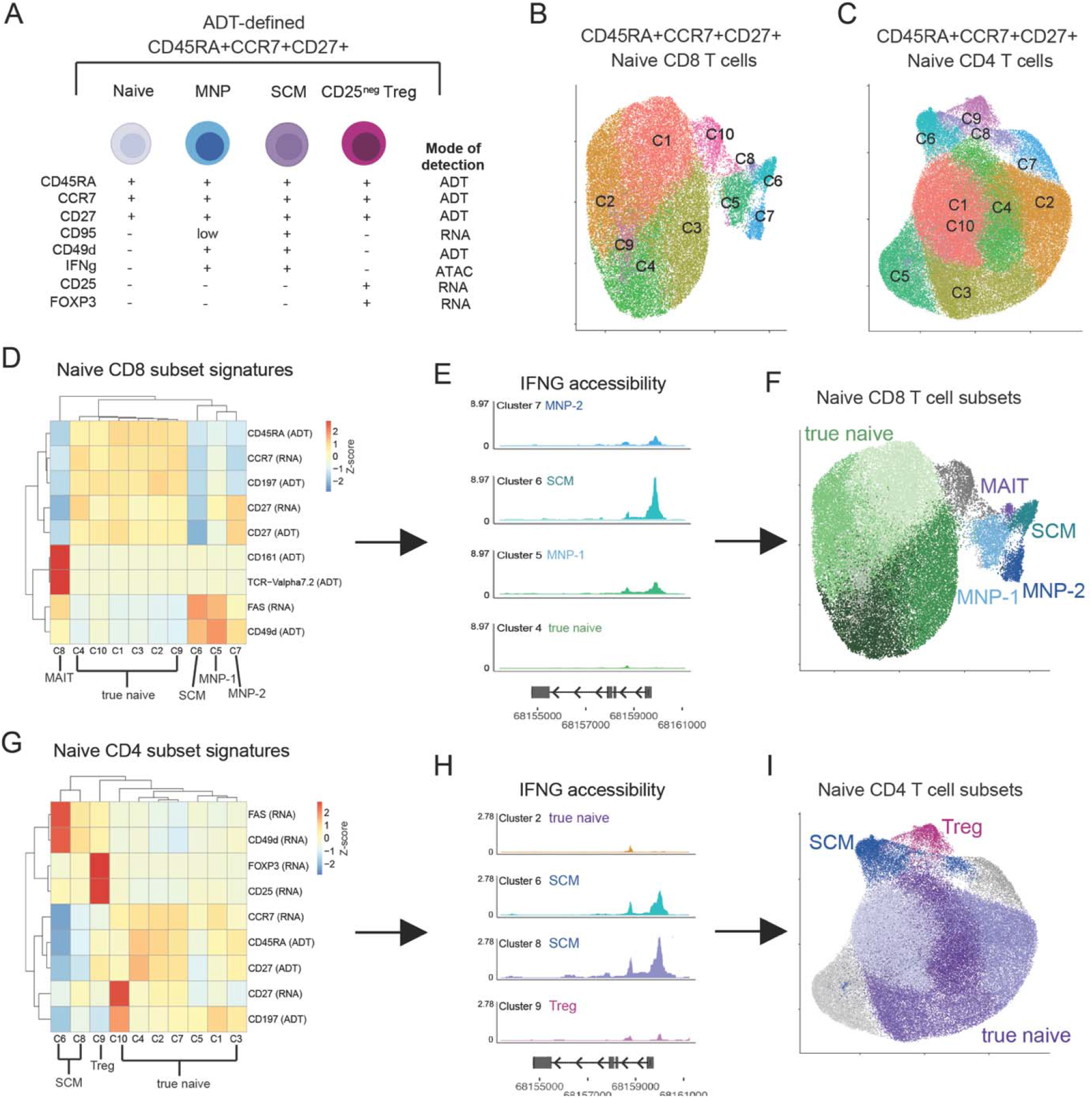
Multiple subsets are present within the phenotypically naive T cell compartment. **(A)** Definition of potential T cell subsets within ADT-defined naive T cells and associated markers with modality of detection. **(B-C)** UMAP representation of 3WNN unsupervised clustering of ADT-based naive **(B)** CD8 and **(C)** CD4 T cells. **(D-E)** Expression of subset markers in naive **(D)** CD8 and **(G)** CD4 T cell clusters. Modality of detection in paratheses. Chromatin accessibility tracks of IFNG gene region on select naive **(E)** CD8 and **(H)** CD4 T cell clusters. **(F)** 3WNN UMAP of naive CD8 T cell subsets, with subsets colored by multi-modal identification, including naive, stem cell-like memory cells (SCM), memory-like naive precursor (MNP-1, MNP-2) and MAIT cells. Gray clusters are not included in subsets as they are considered technical artifacts. **(I)** 3WNN UMAP of naive CD4 T cell subsets, with subsets colored by multi-modal identification, including naive, SCM and CD25^neg^ Tregs. Gray clusters are not included in subsets as they are considered technical artifacts.

After the identification of multiple subsets within (CD45RA^+^CCR7^+^CD27^+^) naive T cells, we next determined the relationship of these subsets to each other and to the overall T cell compartment. To gain initial insight, we first labeled the naive T cell subsets in the context of all T cells in our 3WNN UMAP. (**Figure 3A**) As expected, true naive cells grouped together within the main naive clusters. However, we found that CD8 SCM and CD8 MNP-1 as well as CD4 SCM were grouped with memory T cells in our projection. CD8 SCM clustered closely with effector populations, whereas CD4 SCM spread across multiple memory CD4 T cell populations. Of note, CD8 MNP-1 and CD8 MNP-2 did not group together but instead CD8 MNP-1 grouped with CD8 CM T cells and CD8 MNP-2 grouped with a population of T cells independent of the main central and effector memory populations. CD25^neg^ Tregs closely clustered with CD25^+^CD127^neg^ Tregs. The association of naive T cell subsets with specific memory cell types had little correlation with ADT expression patterns, apart from KLRG1 expression that delineated more effector-like populations (i.e., CD8 TEMRA, CD8 EM2) and CD8 SCM. (**Figure 3B**) However, analyses of RNA and ATAC profiles showed significant molecular similarities between specific naive T cell subsets and other memory T cells. Among CD8 T cells, true naive CD8 T cells grouped separately both by RNA and ATAC, CD8 SCM grouped with TEMRA and EM2, and CD8 MNP-1 cells most closely clustered with CD8 CM1 and CD8 EM1, while CD8 MNP-2 cells had a distinct RNA and ATAC profile from all populations. (**Figure 3C**) Within the naive CD4 T cell subsets, the relationships are somewhat different than in naive CD8 T cells. CD4 SCM cells shared similar molecular profiles as the CD4 CM and CD4 EM1 populations, and the CD25neg Treg CD4 naive subset had profiles most like that of true naive CD4 T cells. (**Figure 3D**) However, the CD25neg Treg CD4 naive subset and the main CD4 Treg population did share some distinct RNA signatures, likely driving their colocalization in 3WNN UMAP projection. (**Figure 3A**) Collectively, these data demonstrate that naive subsets are varied in their transcriptional and epigenetic profiles, with some naive T cell subsets aligning with distinct populations of memory cells.

**Figure 3.**
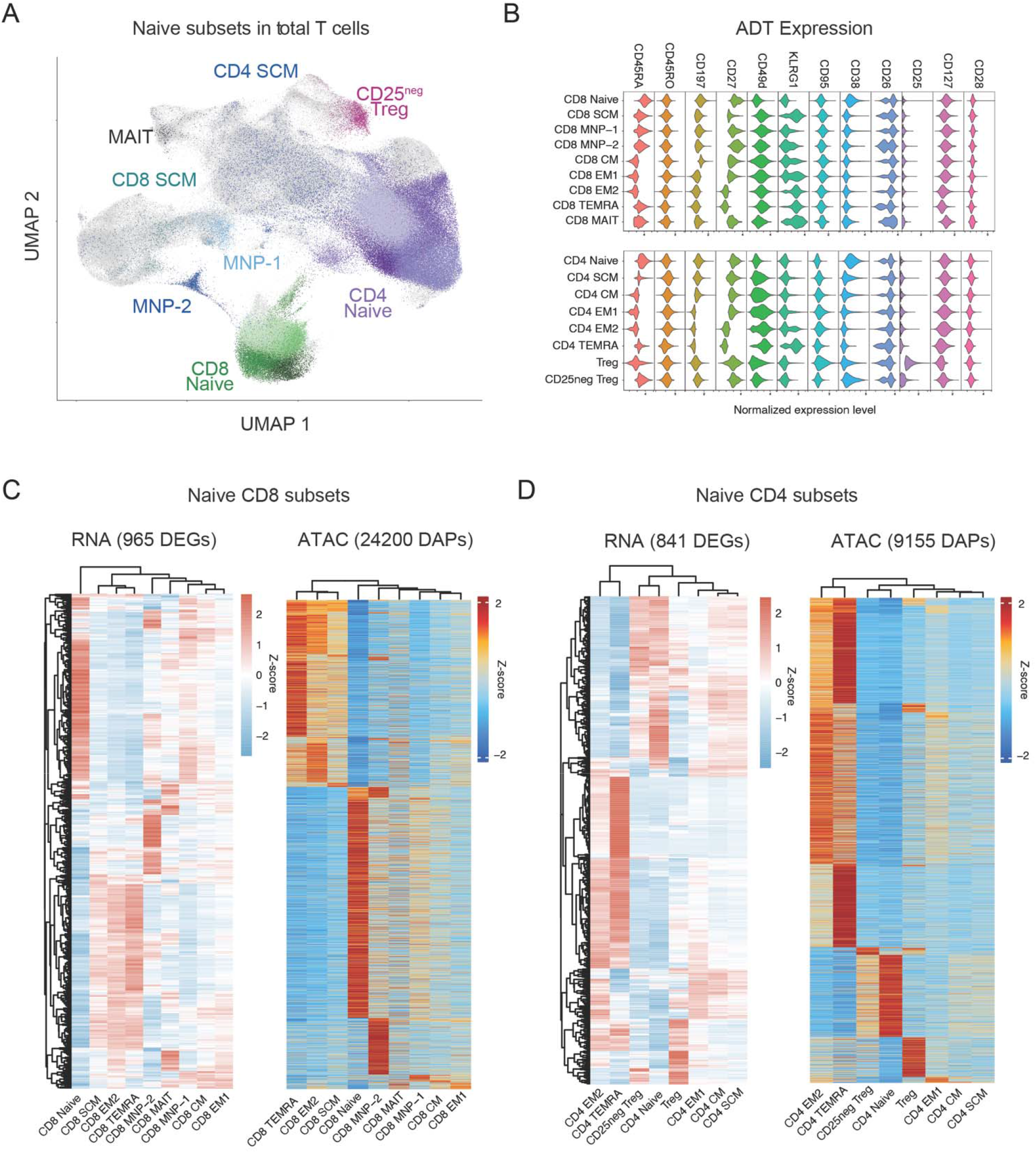
Relationship of transcriptional and epigenetic profiles of naive T cell subsets across the total T cell compartment. **(A)** Tri-modality identified naive CD4 and CD8 T cell subsets overlaid on the 3WNN UMAP of total T cell subsets from all donors. **(B)** Select ADT expression on naive T cell subsets. **(C)** RNA and ATAC profiles of naive CD8 T cell subsets in comparison with ADT-defined memory CD8 T cell populations. **(D)** RNA and ATAC profiles of naive CD4 T cell subsets in comparison with ADT-defined memory CD4 T cell populations.

Naive CD8 T cell subsets were composed of three distinct memory-like subsets: CD8 SCM, CD8 MNP-1 and CD8 MNP-2, that associated with CD8 EM/TEMRA, CD8 CM or no other memory cell respectively. All naive CD8 T cell subsets expressed naive-like transcription and quiescence factors such as LEF1, BACH2 and FOXP1, however each subset also expressed a unique profile of integrins, NK surface receptors, transcription factors and effector molecules. (**Supp Table 2, Supp Fig 5**) Moreover, CD8 SCM and CD8 MNP subsets displayed distinct chromatin accessibility compared with true naive CD8 T cells, with enriched TF motif accessibility for TFs related to effector function such as JUN/FOS, EOMES and TBX21 (Tbet) found in CD8 SCM and CD8 MNP-1. (**Supp Fig 6A-B**) CD8 MNP-2 cells had a distinct TF profile enriched in KLF and SP motifs (**Supp Fig 6C**), suggesting that CD8 SCM and CD8 MNP-1 are more similar to classical CD8 memory cells, whereas CD8 MNP-2 are a non-classical CD8 subset. Thus, we identified multiple types of non-naive cells within the “naive” CD8 T cell compartment that each display unique molecular signatures and are programmed for distinct functional roles.

### Reorganization of memory-like T cell subsets in the naive CD8 T cell compartment with age

In adult aging, the compositional heterogeneity of the “naive” compartment significantly is impacted over time, involving the expansion of CD8+ stem cell memory (SCM) and memory-like naive precursors (MNP) populations in older adults. (*3*, *4*, *19*) However, it is unclear whether compositional changes found in adult aging extend into the naive T cell compartment of children. To determine the impact of age on the composition of naive T cell subsets, we next assessed the frequencies of our tri-modally identified “naive” CD8 T cell subsets in children compared with adults. We found a significant increase in the frequencies of CD8 SCM (1.8% pediatric vs 7% adult; p = 0.02) and CD8 MNP-1 (3% pediatric vs 7.9% adult; p= 0.007) T cell subsets in adults compared with children. (**Figure 4A**) However, the CD8 MNP-2 subset exhibited an inverse correlation with age: CD8 MNP-2 cell frequency was significantly decreased in adults (3.1% pediatric vs 0.8% adult; p = 0.0002). No difference in the frequencies of true naive CD8 T cells was observed. CD8 MAIT cells also showed no significant differences between adults and children. Collectively, naive-like memory CD8 T cells highly expanded from ~8% in children to ~16% in adults. (**Figure 4B**) Thus, aging causes significant alterations in the composition of the naive CD8 T cell compartment through skewing of naive-like memory T cell subset frequencies.

**Figure 4.**
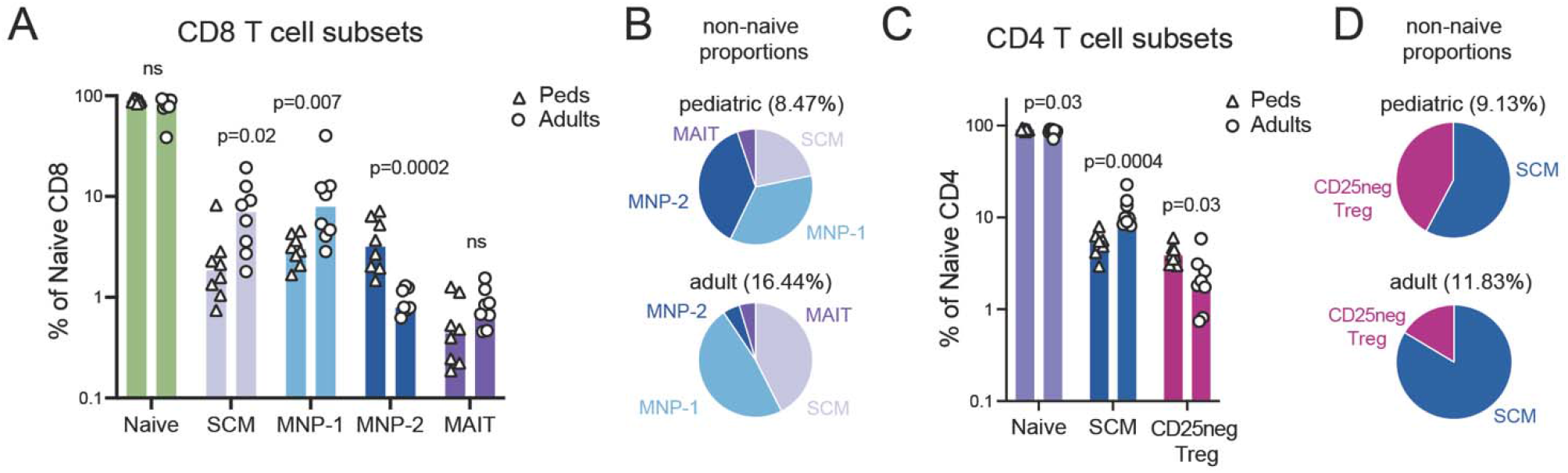
Naive T cell subset frequencies in children compared with adults. **(A)** Frequencies of tri-modally identified subsets within the naive CD8 compartments by age group. P-values determined by Mann-Whitney with Holm-Sidak multiple comparison method. **(B)** Median proportions of the non-naive CD8 subsets in pediatric and adult naive CD8 T cells. Percentages indicate the total frequency of non-naive cells within the compartment. **(C)** Frequencies of tri-modally identified subsets within the naive CD4 compartments by age group. P-values determined by Mann-Whitney with Holm-Sidak multiple comparison method. **(D)** Median proportions of the non-naive CD4 subsets in pediatric and adult naive CD4 T cells. Percentages indicate the total frequency of non-naive cells within the compartment.

We next compared the frequencies of tri-modally identified naive CD4 T cell subsets between children and adults. There was a modest but significant shift in subset distribution with age, with increased CD4 SCM (median 5.3% pediatric vs 9.9% adult; p = 0.004) and decreased CD25^neg^ Tregs (median 3.8% pediatric vs 1.9% adult; p = 0.03) in adults compared with children, respectively. (**Figure 4C**) True naive CD4 T cells had a small decrease in cell frequency across age (median 90.8% pediatric vs 87.2% adult; p = 0.03). Collectively, the age-related shifts in non-naive CD4 T cells were minimal within the overall “naive” compartment, accounting for only a 2.7% increase in adults compared with children. (**Figure 4D**) Together, these data (1) demonstrate the ability of a tri-modal single cell technique to enable detailed resolution of naive T cell subsets, (2) establish that the heterogeneous composition of the “naive” compartment changes across age and (3) reveal that the naive CD8 T cell compartment is more impacted by this compositional shift than the naive CD4 T cell compartment.

### True naive CD4 T cells exist in multiple distinct cellular states

Although we found minimal compositional changes in naive CD4 T cell subsets with age, we previously observed that true naive CD4 T cells could be further divided into 4 major clusters (**Figure 2I, 5A**), indicating there may be separable cellular states within the true naive CD4 T cells. To first understand if these clusters within the true naive CD4 T cells represent distinct states, we examined the global transcriptional (RNA) and epigenetic (ATAC) profiles of each cluster, comparing differentially expressed genes and differentially accessible peaks. (**Figure 5B-5C**) Transcriptionally, we found a total of 186 DEGs between the clusters, with all 4 clusters demonstrating differential patterns of gene expression. (**Figure 5B**) Clusters also demonstrated some overlap in gene expression (e.g., cluster 1 and 3), consistent with these clusters representing cell states and not distinct subsets. Epigenetically, we found 12,117 DAPs across clusters, with all four clusters demonstrating differential ATAC profiles. (**Figure 5C**) Here, we observed that there is a consistent set of peaks opening and a set closing stepwise across the clusters, indicative of transitioning molecular states.

**Figure 5.**
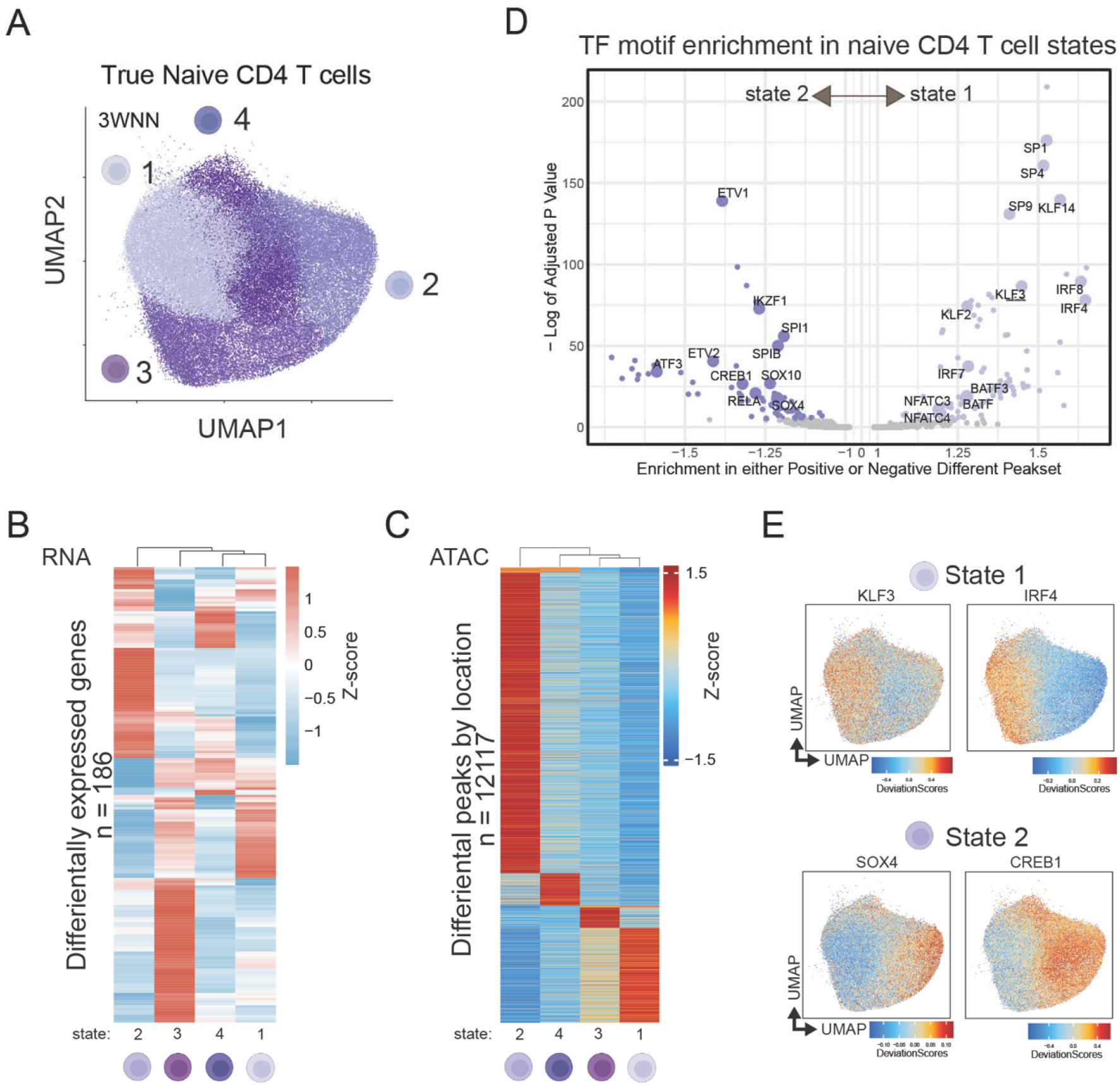
Distinct cell states within the true naive CD4 T cell compartment. **A)** 3WNN UMAP with leiden clustering of true naive CD4 T cells. **(B)** Heatmap of marker genes for the true naive CD4 T cell clusters. Seurat’s FindAllMarkers (parameters logfc.threshold=0.25, pval<0.05) **(C)** Heatmap of marker peaks for true naive CD4 T cell clusters. ArchR’s getMarkerFeatures (parameters maxCells=10000, FDR <= 0.1, Log2FC >= 0.5 **(D)** Transcription factor motif enrichment based on differentially accessible peaks between state 1 and 2. **(E)** 3WNN UMAP of true naive CD4 T cells with ChromVar motif deviations for selected state-enriched transcription factors.

Clusters 1 and 2 demonstrated the most divergence in chromatin landscape, potentially implicating these clusters as functionally distinct cell states. We performed transcription factor motif (ATAC analysis) enrichment comparing these clusters, and found that clusters 1 and 2 displayed differential TF binding motif accessibility. (**Figure 5D**) Cluster 1 was preferentially biased towards chromatin accessibility in regions with TFs related to activation (e.g., KLFs, SP1, NFAT) and cytokine signaling (e.g., IRFs). Conversely, cluster 2 had TF motif accessibility associated with NFKB signaling (e.g., RELB, CREB1), TGFβ signaling (e.g., SOX4) and differentiation (e.g., IKZF1). (**Figure 5E-F**) Moreover, we found that transcriptional changes in these clusters were linked with epigenetic changes, with SOX4 demonstrating essentially no chromatin accessibility in state 1, consistent with virtually no gene expression (**Sup Fig 7**). Collectively, these data reveal that true naive CD4 T cells exist in multiple cell states.

### Distinct molecular programming of pediatric and adult naive CD4 T cells is driven by alterations in cellular state

Our original global analyses demonstrated major transcriptional and epigenetic changes between naive CD4 T cells from children and older adults. (**Figure 1**) We hypothesized that these age-related changes were driven by the composition of the cell states within true naive CD4 T cells. We compared the distribution and frequencies of the 4 cell states of true naive CD4 T cells between children and adults, and found true naive CD4 T cells from adults were significantly skewed towards cluster 1 (12.7% pediatric vs 54.7% adult, median; p = 0.0002). (**Figure 6A-B**) Conversely, true naive CD4 T cells from children were much more evenly distributed across the clusters, with cluster 2 making up a significantly higher proportion of the pediatric naive population than that of adults (36.5% pediatric vs 5.5% adult, median; p = 0.0019). Cluster 4 also displayed a pediatric bias (24.2% pediatric vs 2.5% adult, median; p = 0.0002). Cluster 3 showed no significant distribution differences between children and adults (18.6% pediatric vs 24.1% adult, median; p = 0.2345). Pseudo-bulk expression analysis of true naive CD4 T cells similarly revealed that adults had higher expression of cluster 1 representative gene STAT4 whereas children had higher expression of cluster 2 and 4 representative genes SOX4 and TGFBR2, respectively. (**Figure 6C**) Of note, increased accessibility around BATF and IRF4 motifs in the adult-enriched cluster 1 (**Figure 5D**) is consistent with previous epigenetic comparisons between young and older adults naive CD4 T cells. (*20*) Together, these data demonstrate that age alters the distribution of true naive CD4 T cells across distinct cell states.

**Figure 6.**
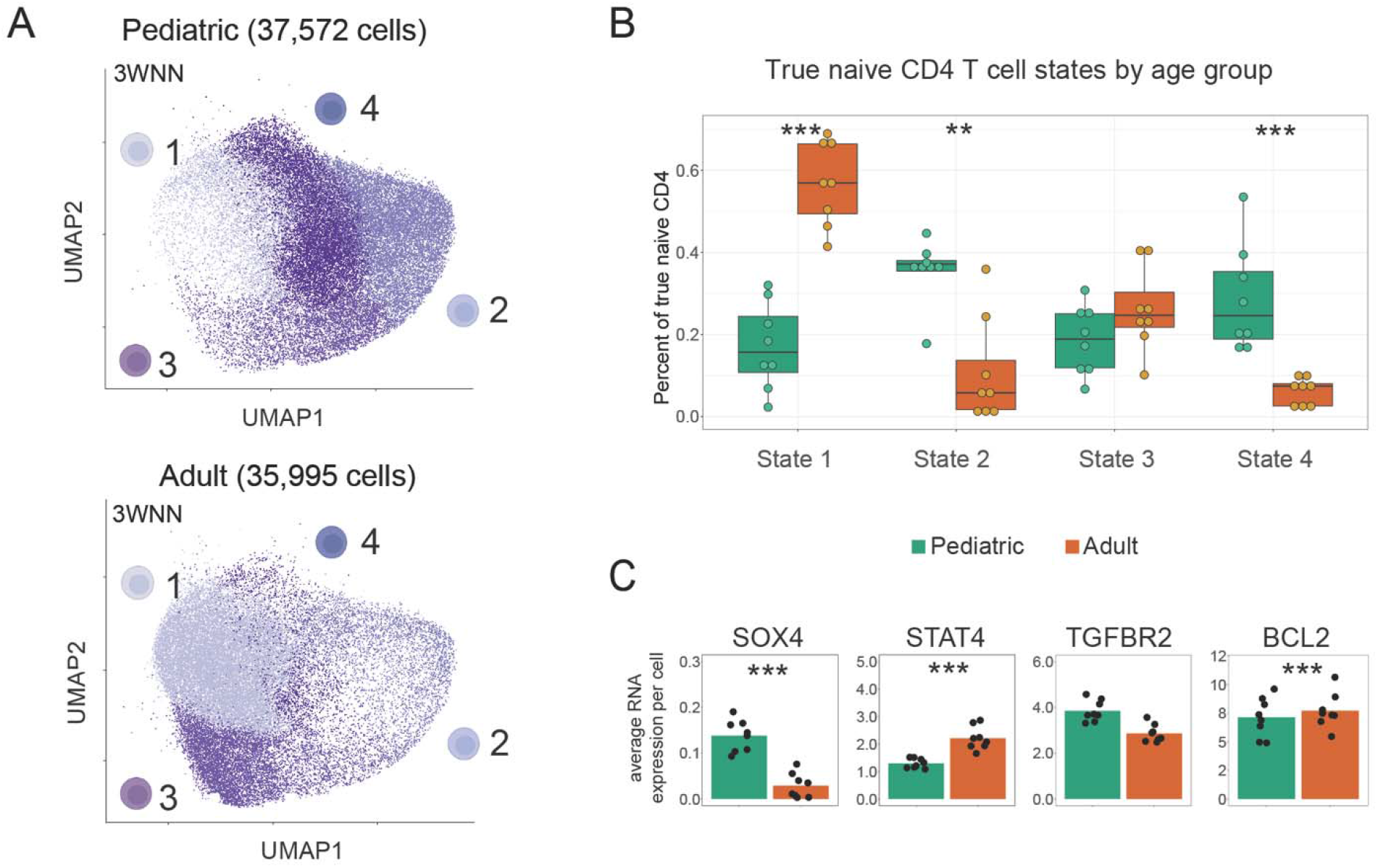
Shift in naive CD4 T cell states between children and adults. **(A)** 3WNN UMAP projection of true naive CD4 T cell clusters by age group. **(B)** Distribution of true naive CD4 T cells within the 4 identified cell cluster by age group. Green = pediatric, orange = adult. Mann-Whitney test. **(C)** Pseudo-bulk expression of select cluster-specific genes within true naive CD4 T cells separated by age. Mann-Whitney test. ** p < 0.01, *** p < 0.001

To further confirm if the shifts in cellular state of true naive CD4 T cells between children and older adults span across adulthood, we performed scRNA-seq on PBMCs from children (n=8, 11-13 yrs), young adults (n=8, 25-35 yrs) and older adults (n=8, 55-65 yrs). We then used Seurat’s reference-based RNA cell label transfer methods to define naive CD4 T cells within the scRNA-seq datasets. When applied to our TEA-seq dataset, we found a 91% agreement between the Seurat naive CD4 T cell labels and our true ADT-based naive CD4 T cell designations, suggesting that Seurat’s label transfer is a reliable method for identifying most, but not all, true naive CD4 T cells when cell surface protein measurements are unavailable. (**Supp Figure 7**)

Naive CD4 T cells were first analyzed for overall transcriptional similarity. Notably, naive CD4 T cells from children grouped separately from those of both young adults and older adults. (**Figure 7A**). We then examined gene signatures that distinguished naive CD4 T cell states (**Figure 5, Supp Table 3**) and found that both young and older adult naive CD4 T cells exhibited similar gene expression, whereas pediatric naive CD4 T cells were more distinct, favoring genes found in cell state 2 (e.g., TOX, IKZF2, PLCB1, SOX4). (**Figure 7B**) Both young adults and older adults displayed little expression of these genes. Naive CD4 T cells from older adults had the highest expression of cell state 1 signature genes (e.g. TSHZ2, CPQ, STAT4). These genes showed increased expression across increasing age categories. Thus, true naive CD4 T cells are in a transcriptionally distinct states in children compared with adults.

**Figure 7.**
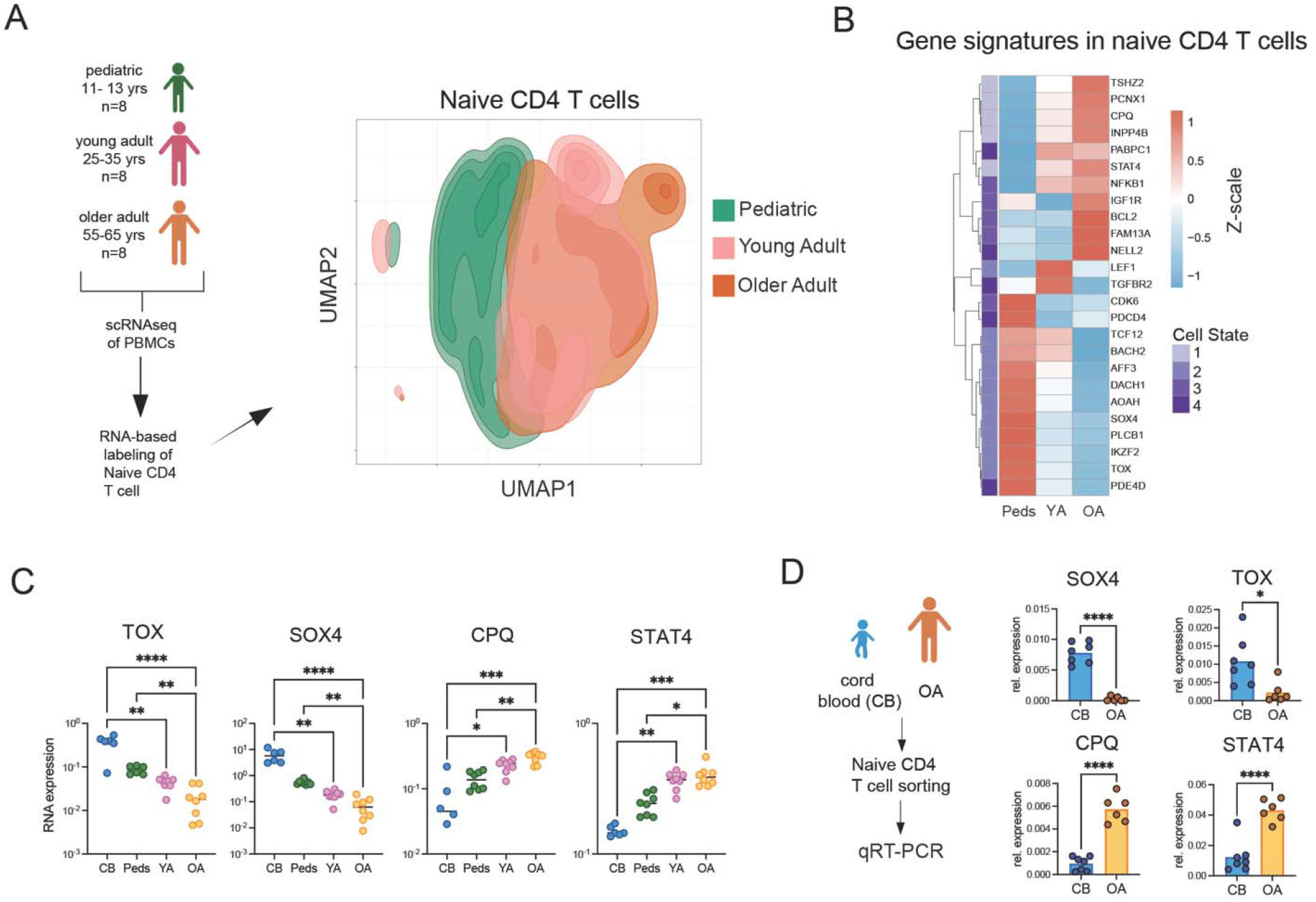
Unique transcriptional profiles of naive CD4 T cells across aging spectrum. **(A)** RNA-based UMAP of naive CD4 T cells (71,034 cells) from scRNA-seq data from children, young adults and older adults (n = 8 subjects per age group). **(B)** Expression of select DEGs from each cell state of RNA-predicted naive CD4 T cells across age groups. **(C)** Average expression of cell state marker genes in naive CD4 T cells from scRNA-seq datasets, including an external cord blood dataset. (*21*, *22*) **(D)** Gene expression of SOX4, TOX, CPQ and STAT4 in bulk sorted naive CD4 T cells (CD3+CD4+CD8-CD45RA+CCR7+CD27+CD95-) from newborn cord blood (n = 7 donors) and older adult peripheral blood (n = 6 donors) as determined by qRT-PCR. Mann-Whitney test. * p < 0.05, ** p < 0.01, *** p < 0.001, **** p < 0.0001

The contrast in pediatric and adult naive CD4 T cell transcriptional profiles suggests that naive CD4 T cells produced early in life have unique molecular features compared to those developed or maintained in adulthood. To see if this trend continued to earlier points in human development, we next asked whether naive CD4 T cells in cord blood from newborn infants also demonstrated the same skew in naive CD4 T cell gene expression as seen in pediatric subjects. Utilizing the same approach as described above on two publicly available scRNA-seq datasets from cord blood (*21*, *22*), we found that cord blood naive CD4 T cells had expression patterns similar to children, with high expression of SOX4 and TOX and conversely low expression of CPQ and STAT4. (**Figure 7C**) To verify that this transcription signature is specific to naive CD4 T cells developing early in life, we then performed RT-qPCR on bulk naive CD4 T cells from a new cohort of cord blood (n=7) and older adult peripheral blood (n=6, >60 yrs) using fluorescence-assisted cell sorting (FACS) based on the following protein expression pattern: CD4^+^CD8^−^CD45RA^+^CCR7^+^CD27^+^CD95^−^. Consistent with our TEA-seq and scRNA-seq experiments, cord blood naive CD4 T cells demonstrated significantly higher expression of TOX and SOX4, and decreased expression of CPQ and STAT4 compared with older adults. (**Figure 7D**) Indeed, SOX4 was essentially undetectable in adult naive CD4 T cells. Together, these data demonstrate that true naive CD4 T cells shift towards transcriptionally separable states across lifespan, and that the pediatric-specific transcriptional programming arises early in life.

## DISCUSSION

Aging has a profound impact on naive T cells. However, our understanding of the complexity of this impact across lifespan has been limited. Here, we utilize TEA-seq to simultaneously interrogate the cellular and molecular heterogeneity of naive T cells in children and adults. We established that age causes significant changes in composition as well as the transcriptome and epigenome of naive T cells, in contrast to CMV infection status that preferentially impacts effector populations. Detailed interrogation of naive T cell subsets revealed that the compositional changes in the naive CD8 T cell compartment drive overall molecular alterations, with a unique memory-like subset preferentially lost in adults. We discovered that age-related molecular differences in naive CD4 T cells were driven by preferential shifts in cellular state of the true naive CD4 T cell subset. Collectively, these changes may contribute to differential responses to infection, vaccination and therapeutic treatments in children and older adults.

One of the most consistent features of adult aging is the loss of naive CD8 T cells. However, a few other changes have been noted in this compartment with age: 1) memory cell infiltration, 2) pseudo-differentiation and 3) clonal expansion. (*23*) Although we were currently unable to assess clonal expansion in this study, we directly interrogated memory infiltration and pseudo-differentiation in the naive CD8 T cell compartment of adults versus children. We delineated multiple cell subsets within the *resting* naive CD8 T cells – including true naive CD8, CD8 SCM, CD8 MAIT and two populations of memory-like naive CD8 precursors. Consistent with previous publications on adult aging, there was significant expansion of CD8 SCM and CD8 MNP-1 in adults compared with children, suggesting that these populations are indeed developed from antigen-specific responses over the course of a lifespan. (*2*) These two memory-like populations within the naive CD8 T cell compartment demonstrated transcription and epigenetic signatures similar to those found in previous bulk studies of naive CD8 T cell aging. (*5*) These data are also consistent with a recent RNA-seq study that found minimal evidence for pseudo-differentiation of naive CD8 T cells in young compared with older adults. (*24*) Collectively, our results implicate the infiltration of specific memory-like subsets into the “naive” compartment as a primary driver of naive CD8 T cell aging.

Unlike naive CD8 T cells, the naive CD4 T cell compartment is considered relatively resistant to aging, showing less memory infiltration and pseudo-differentiation in adult aging. (*20*, *25*) However, using our multi-modal classification strategy, we find that the true naive CD4 T cells exist in multiple distinct cellular states and that there is a significant age-related bias for which states naive CD4 T cells are maintained in. Pediatric naive CD4 T cells were primarily present in a cellular state indicative of quiescence whereas adult naive CD4 T cells were biased towards a more activated state. The relative subtlety of the change (i.e., in the absence of any major changes in surface homing markers expression, etc.) is similar to a newer concept in the stem cell aging field in which quiescent stem cells move into a primed quiescent state upon bystander exposure to the aging microenvironment. (*26*) In the context of T cell aging, multiple studies in animal models reveal increased fibrosis in lymph nodes with age is associated with impaired T cell homeostasis. (*27*, *28*) Thus, investigation into the crosstalk between naive T cell state and the microenvironment is of great interest.

In the context of stem cell biology, primed quiescent stem cells gain the ability to rapidly respond to tissue injury, however this comes at a cost to their pluripotency. (*26*) The finding that naive CD4 T cells from adults are more likely to be in a cell state that is primed or activated suggests that the differentiation capacity of these cells may also be impacted. Functional studies on cord blood naive T cells previously demonstrated that these cells are not intrinsically predisposed to be better effector cells, but under inflammatory conditions will produce greater amounts of IFNγ than adult naive CD4 T cells. (*29*) This suggests that the cellular state of naive CD4 T cells early in life is primed to respond more rapidly to external stimulation than in adults. This is similar to a concept in mouse naive CD8 T cell aging, in which naive CD8 T cells from infant mice were epigenetically pre-programmed to favor effector over memory formation. (*6*, *7*)

In our analysis of true naive CD4 T cells, we specifically identified SRY-box transcription factor 4 (SOX4) as signature TF of the pediatric-enriched naive CD4 T cell state. SOX4 expression in T cells is upregulated by TGF-beta signaling and inhibited by IL-2, (*30*, *31*) and plays a role in the regulation of naive T cell quiescence. (*32*) Additionally, SOX4 activity induces the expression of miRNA-181a and leads to enhanced T cell receptor signaling. (*33*) In adults, the downregulation of miRNA-181a with age causes reduced TCR signaling in naive T cells. (*25*, *34*, *35*) This downregulation of TCR signaling is also in tandem with the preferential development into memory precursors over effector phenotypes. (*36*) Tonic TCR signaling strength has also been implicated in shifts in functionality of naive T cells (*37*) and could influence regulation of cell state over the course of a lifetime. Whether the loss of SOX4 may be a pioneer transcription factor for effector differentiation or an upstream driver of altered outcomes to antigen-specific responses with aging remains to be determined.

Collectively, these studies demonstrate that phenotypically naive T cells are highly heterogeneous and that the CD8 and CD4 naive T cell compartments are impacted differently by age - displaying major changes in composition and cell programming with age, respectively. Although solely gleaned from female volunteers, these findings have significant translational implications for vaccine development, highlighting that naive T cells in children are not the same as in adults, and their response to the same stimulation likely differs. It also demonstrates that TEA-seq could be easily applied to enhance our understanding of T cell subsets in many auto-immune and/or inflammatory disease states, such as rheumatoid arthritis, HIV and obesity, and can be used to establish molecular drivers of T cell dysfunction as potential novel targets of therapeutic intervention.

## MATERIALS AND METHODS

### Adult and Pediatric Cohorts

*Adult*: Healthy 55-65 year old adult subjects were recruited from the greater Seattle area as part of the Sound Life project, a protocol approved by the Institutional Review Board (IRB) of the Benaroya Research Institute. Patients were excluded from enrollment if they had a history of chronic disease, autoimmune disease, severe allergy, or chronic infection. *Pediatric:* Healthy 11-13 year old pediatric subjects were recruited from the greater Philadelphia area under a protocol approved by the IRB of the Children’s Hospital of Philadelphia. Patients were excluded from enrollment if they had a history of immune deficiency, fever or antibiotic usage within the month prior to sample collection, chronic medication usage, or BMI more than 2 standard deviations above or below the mean for their age. We selected only female donors, to limit sex-based differences. Half the pediatric and adult donors were CMV-positive, based on testing in a CLIA-approved lab. All samples were collected, processed to PBMCs using a Ficoll-based approach and frozen in FBS with 10% DMSO within 4 hours of blood draw. Cord and peripheral blood samples for follow-up studies were purchased from Bloodworks Northwest (Seattle, WA) through protocols approved by the Bloodworks Northwest and internal Allen Institute IRBs.

### TEA-seq

TEA-seq library preparation was performed as described previously (*13*), with the addition of Cell Hashing (*38*) to allow for sample multiplexing and limit well-to-well batch effects. In brief, samples were thawed and processed across three batches, with each batch containing a common PBMC control. Antibody staining for Cell Hashing and cell sorting was performed simultaneously on 2 × 10^6^ cells from each sample. Each sample was incubated with a sample-specific barcoded TotalSeq-A antibody, anti-CD45 antibody and anti-CD3 antibody, then pooled by T cell proportions previously determined by flow cytometry, targeting 800,000 T cells for each donor sample and 200,000 T cells for the control, and sorted on a BD FACSAria Fusion. T cells were sorted as live, single CD45^+^CD3^+^ cells. 2 × 10^6^ sorted T cells were then used for the library preparation. A panel of 55 target-specific barcoded oligo-conjugated antibodies (BioLegend TotalSeq-A) was used for these studies (**Supp Table 4**). Individual ATAC, RNA, and ADT libraries were prepared, sequenced and processed as described previously.(*13*)

### TEA-seq Data Pre-processing

ADT and HTO count matrices were generated using BarCounter v1.0 (*39*). The RNA count matrix, and ADT count matrix were then combined into a single Seurat object. Cells were selected based on the following cutoffs: > 250 genes/cell, > 500 RNA UMIs/cell, < 10,000 ADT UMIs/cell, < 35% mitochondrial reads, < 20,000 RNA UMIs/cell. Normalization, feature selection and scaling were performed on the RNA matrix (Seurat SCTransform function, default settings), followed by PCA (Seurat RunPCA function, default settings). A UMAP projection was generated (Seurat RunUMAP, with parameter dims = 1:30) and clustering was performed (Seurat FindNeighbors, with parameter dims = 1:30, followed by Seurat FindClusters, with parameter resolution = 0.5). We used the Seurat Multimodal Reference Dataset for PBMCs (available from the Satija lab (*15*)) to perform label transfer on the dataset using the functions described in the Seurat V4 Vignettes (Seurat FindTransferAnchors, followed by Seurat TransferData). Two clusters were identified to be non-T cells and were excluded from downstream analysis. Sample specific transcripts, AC105402.3 and MTRNR2L8, were identified and removed prior to further downstream RNA analysis.

### ADT-based Cell Type Identification

We used CD4, CD8, CD197, CD27 and CD45RA ADT markers to identify T cell subsets. For subset identification, each of the three batches were separated into its own Seurat object before analysis to account for differences in sequencing depth and average ADT UMIs/cell. ADTs were normalized and cells were identified based on the markers outlined in Figure 1 and **Supplemental Table 1** using Boolean gating.

### ADT, RNA and ATAC Label Transfers

RNA-based label transfer was performed using single positive T cell subsets from the Seurat reference described above using Seurat functions FindTransferAnchors and TransferData. Label transfer from ATAC data was performed using the same reference, based on ArchR (version 1.0.2) documentation (https://archrproject.com). (*40*) A first round of unconstrained integration was performed, and cells were labeled based on the Seurat L1 cell types. The second round of labeling then used the constrained approach to transfer the L2 cell types based on the results of the L1 integration. To directly compare the results from both RNA and ATAC label transfer with our ADT-defined populations, select cell types were merged manually.

### TEA-seq T cell Subsets Analyses

<u>*3WNN Clustering.*</u> We performed PCA on both RNA and ADT count matrices and then corrected for any potential batch effects using Harmony (https://github.com/immunogenomics/harmony). (*41*) For ATAC, an LSI embedding was calculated in ArchR (ArchR addIterativeLSI function, with parameter varFeatures = 75000), and batch correction performed (ArchR addHarmony function, with parameter groupBy = ‘batch_id’). The resulting harmony corrected LSI embedding was transferred over to the Seurat object for 3-way weighted nearest neighbors (3WNN) integration and clustering on all harmony corrected PCAs/LSI (Seurat FindMultiModalNeighbors function, with parameter dims.list = list(1:25, 1:20, 1:29) for RNA, ADT, ATAC respectively). <u>*RNA modality analysis.*</u> Differentially expressed gene (DEG) analysis was performed with the hurdle model implemented in the MAST package (*42*). The p-values were adjusted for multiple comparisons using the Benjamini and Hochberg (BH) method. (*43*) Adjusted p-values < 0.05 and logFC > 0.1 were considered significant. <u>*ATAC modality Analysis.*</u> Arrow files generated by the TEA-seq processing pipeline previously described were filtered to retain single-positive T cells. These files were then merged into a single ArchR project for Iterative LSI, Clustering, Group Coverage computation, Reproducible Peak Set annotation, Motif Enrichment and ChromVar Deviations Enrichment according to the ArchR documentation. The Peak Matrix was used to identify differentially accessible peaks between groups. The identified peaks were used in a motif enrichment analysis (ArchR peakAnnoEnrichment function, with cutoffs FDR ≤ 0.1 & Log_2_FC ≥ 0.5). Chromatin accessibility tracks and ChromVar motif enrichment plots were generated by scripts in this repository (https://github.com/aifimmunology/scATAC_Supplements).

### DEG Pathway Enrichment Analysis

Pathway enrichment analysis was performed with Gene Set Enrichment Analysis (GSEA) (*44*) implemented in the FGSEA package (*45*) to compare between the adult and pediatric subjects and by their CMV status, respectively. A custom collection of genesets that included the Hallmark v7.2 genesets, KEGG v7.2 and Reactome v7.2 from the Molecular Signatures Database (MSigDB, v4.0) was used as the pathway database, as previously described. (*46*) The pathway enrichment p-values were adjusted using the BH method and pathways with adjusted p-values < 0.05 were considered significantly enriched.

### Transcription Factor Motif analysis

For each ADT-labeled cell type, age group (i.e., pediatric versus adult) or CMV infection status were compared to identify differentially accessible peaks (ArchR getMarkerFeatures funtion) using the maximum number of overlapping cells between these groups. Motif enrichment (ArchR peakAnnoEnrichment function) was then performed using the resulting differentially accessible peaks with an FDR cutoff of ≤ 0.1 and Log_2_FC cutoff ≥ 0.5. Motifs for each cell type were then further filtered by mlog10Padj > 5 cutoff and found differentially expressed in at least 6 of the cell types. No enriched motifs were detected based on CMV status, so no plots were generated for visualization.

### Naive CD4 and CD8 T cell Sub-Analysis

We performed 3WNN clustering as described above for ADT-identified CD4^+^ and CD8^+^ naive T cells separately. Leiden clusters were then identified at multiple resolutions by varying the resolution parameter of the Seurat FindClusters function from 0.1 to 0.8, and were visualized using the Clustree package (*47*) (https://github.com/lazappi/clustree) to identify the optimal resolution. To maintain robust comparisons, clusters with fewer than 650 cells, as well as clusters driven by expression of a single gene, were removed from downstream analysis. Marker genes for each cluster were then identified using Seurat’s FindAllMarkers function. ATAC analysis was performed on the same separated populations using the same approach described above in ArchR.

### Flow Cytometry

To assess T cell subset frequencies, PBMCs were analyzed using a 25-color T cell phenotyping flow cytometry panel (**Supp Table 4**), using the standardized method previously published. (*48*) Cells were analyzed on a 5 laser Cytek Aurora spectral flow cytometer. Spectral unmixing was calculated with pre-recorded reference controls using Cytek SpectroFlo software (Version 2.0.2). Cell types were quantified by traditional bivariate gating analysis performed with FlowJo cytometry software (Version 10.8)

### scRNA-seq

Single cell RNA sequencing was performed on PBMCs as previously described. (*48*) In brief, scRNA-seq libraries were generated using a modified 10x genomics chromium 3’ single cell gene expression assay with Cell Hashing. 8 donors were pooled per library, with the addition of a common batch control sample in each library. Libraries were sequenced on a Illuminia Novaseq platform. Hashed 10x Genomics scRNA-seq data processing was carried out using BarWare (*39*) to generate sample-specific output files. *scRNA-seq Analysis.* Count matrices from each sample were merged into age-specific Seurat objects, followed by normalization, feature selection, scaling, PCA, UMAP embedding, and clustering, as described above. Label transfer from the T cell fraction of the PBMC Seurat reference was performed as described above for each age-specific dataset. Following label transfer, all objects were merged into a single dataset. Cells identified as CD4 Naive with a prediction score greater than 0.7 were retained for downstream analysis. We then averaged the expression on each cell in each age group (Seurat AverageExpression function, group.by = ‘age’) for DEGs identified in our TEA-seq analysis for use in visualization.

### Naive CD4 T cell Sorting

T cells were directly isolated from peripheral or cord blood using the RosetteSep Human T-cell Enrichment Cocktail according to the manufacturer’s protocol (Stem Cell Technologies cat#15061). T cells were cryopreserved in 90% fetal bovine serum+10% DMSO and stored in vapor phase liquid nitrogen following isolation. Cryopreserved T Cells were rapidly thawed and stained with the sorting antibody panel in **Supp Table 4**. Naive CD4 T cells were sorted using the Melody Sorter (BD Biosciences) according to the following phenotype: Live, single, CD3^+^CD8^−^ CD4^+^CCR7^+^CD45RA^+^CD27^+^CD95^−^ cells. 500,000 cells per sample were then pelleted and snap frozen in dry ice/ethanol for RNA isolation.

### RNA extraction and qPCR

Total RNA was isolated using the RNeasy Mini Plus (Qiagen 74034) or Micro Plus Kit (Qiagen 74134) per manufacturer protocol. cDNA was generated using the Supercript IV VILO Master Mix (Invitrogen 11755050). For qPCR, Taqman probes sets (assay IDs: RPLP0 Hs999999902_m1; TOX Hs01049519_m1, SOX4 Hs04987498_s1, CPQ Hs01550609_m1 and STAT4 Hs01028017_m1) using Taqman Fast Advanced Master Mix on the BioRad CFX96 Real-Time Instrument. All genes were normalized to housekeeping gene, RPLP0 and expression compared using the 2^(-dCt)^ method.

## Supporting information

SupTable2

SupTable3

## Supplementary Materials

Supplemental Figure 1. Flow cytometry hand-gating of T cell subsets

Supplemental Figure 2. Comparison of ADT-defined T cell subsets in flow cytometry and age.

Supplemental Figure 3. Cross-modal analysis of T cell subsets.

Supplemental Figure 4. Frequencies of T cell subsets by age and CMV infection status.

Supplemental Figure 5. Select RNA profile of naive CD8 T cell subsets.

Supplemental Figure 6. TF motif enrichment across naive CD8 T cell subsets.

Supplemental Figure 7. Chromatin accessibility tracks of true naive CD4 T cell states

Supplemental Figure 8. Confusion plot comparison

Supplemental Table 1: Cell Subset Markers for ADT-based Identification

Supplemental Table 2. Differentially expressed genes between Naive CD8 T cell subsets.

Supplemental Table 3. Differentially expressed genes between true naive CD4 T cell states.

Supplemental Table 4. Antibody List

## Acknowledgements

We thank the study participants and the clinical research teams at University of Pennsylvania, CHOP and Benaroya Research Institute for their dedication to this project. We thank the Allen Institute founder, Paul G. Allen, for his vision, encouragement, and support. We also thank all the members of the Allen Institute for Immunology, in particular the facilities and operations teams who helped maintain the productive research environment, as well as the Human Immune System Explorer (HISE) software development team for their constant support. Research reported in this publication was supported by the National Institute on Aging of the National Institutes of Health under Award Number K01AG068373 and by the Allen Institute for Immunology. This research was conducted while Claire E. Gustafson was a Diamond/AFAR award recipient. The content is solely the responsibility of the authors and does not necessarily represent the official views of the National Institutes of Health.

## Funding

National Institutes of Health grant K01AG068373 (CEG)

American Federation for Aging Research (CEG)

Allen Institute for Immunology

## Author contributions

Conceptualization:CEG,ZT,ES,JR,LTG,GLS,AKS,CS,JHB,XJL,TRT,EJW,TFB,LAV,SHH,PJS

Methodology:ZT,ES,KH,MLP,LYO,ATH,CRR,VH,MDAW,PCG,JR,JRG,SM,LTG,SVV,GLS,AKS,TRT,TFB, LAV,SHH,PJS

Investigation:CEG,ZT,ZH,KH,MLP,LYO,ATH,CRR,VH,MDAW,PCG,JR,JRG,SM,JD,CJJ,ARG,XJL,TRT,E JW,LAV,SHH,PJS

Visualization: CEG,ZT,ZH,KH,MLP,LYO,TRT

Funding acquisition: LAB,TRT,TFB,PJS

Project administration: JD,CJJ,ARG,LAB,CS,JHB,EJW,PJS

Supervision: CEG,ZT,LTG,SVV,GLS,TRT,PJS

Writing – original draft: CEG,ZT,ZH,PJS

Writing – review & editing:

CEG,ZT,ZH,JR,LTG,SVV,GLS,AKS,CS,JHB,TRT,EJW,TFB,LAV,SHH,PJS

## Data and Materials Availability

Raw data will be deposited in the NCBI Database of Genotypes and Phenotypes (dbGaP, Accession ID: TBD) for controlled access upon peer reviewed publication. Processed data will be deposited in the NCBI Gene Expression Omnibus database (GEO, Series Accession ID: GSE214546). Code used for analysis and figure generation in this manuscript will be available on Github upon peer reviewed publication. External cord blood scRNA-seq datasets are from GEO database (IDs: GSE157007 and GSE165193). (*21*, *22*)

