## Supplemental Materials for "Distinct Heterogeneity in the Naive T cell Compartments of Children and Adults"

**Supplementary Materials**

Supplemental Figure 1. Flow cytometry hand-gating of T cell subsets

Supplemental Figure 2. Comparison of ADT-defined T cell subsets in flow cytometry and age.

Supplemental Figure 3. Cross-modal analysis of T cell subsets.

Supplemental Figure 4. Frequencies of T cell subsets by age and CMV infection status. Supplemental Figure 5. Select RNA profile of naïve CD8 T cell subsets.

Supplemental Figure 6. TF motif enrichment across naïve CD8 T cell subsets.

Supplemental Figure 7. Chromatin accessibility tracks of true naïve CD4 T cell states

Supplemental Figure 8. Confusion plot comparison

Supplemental Table 1: Cell Subset Markers for ADT-based Identification

Supplemental Table 2. Differentially expressed genes between Naïve CD8 T cell

subsets.

Supplemental Table 3. Differentially expressed genes between true naïve CD4 T

cell states.

Supplemental Table 4. Antibody List

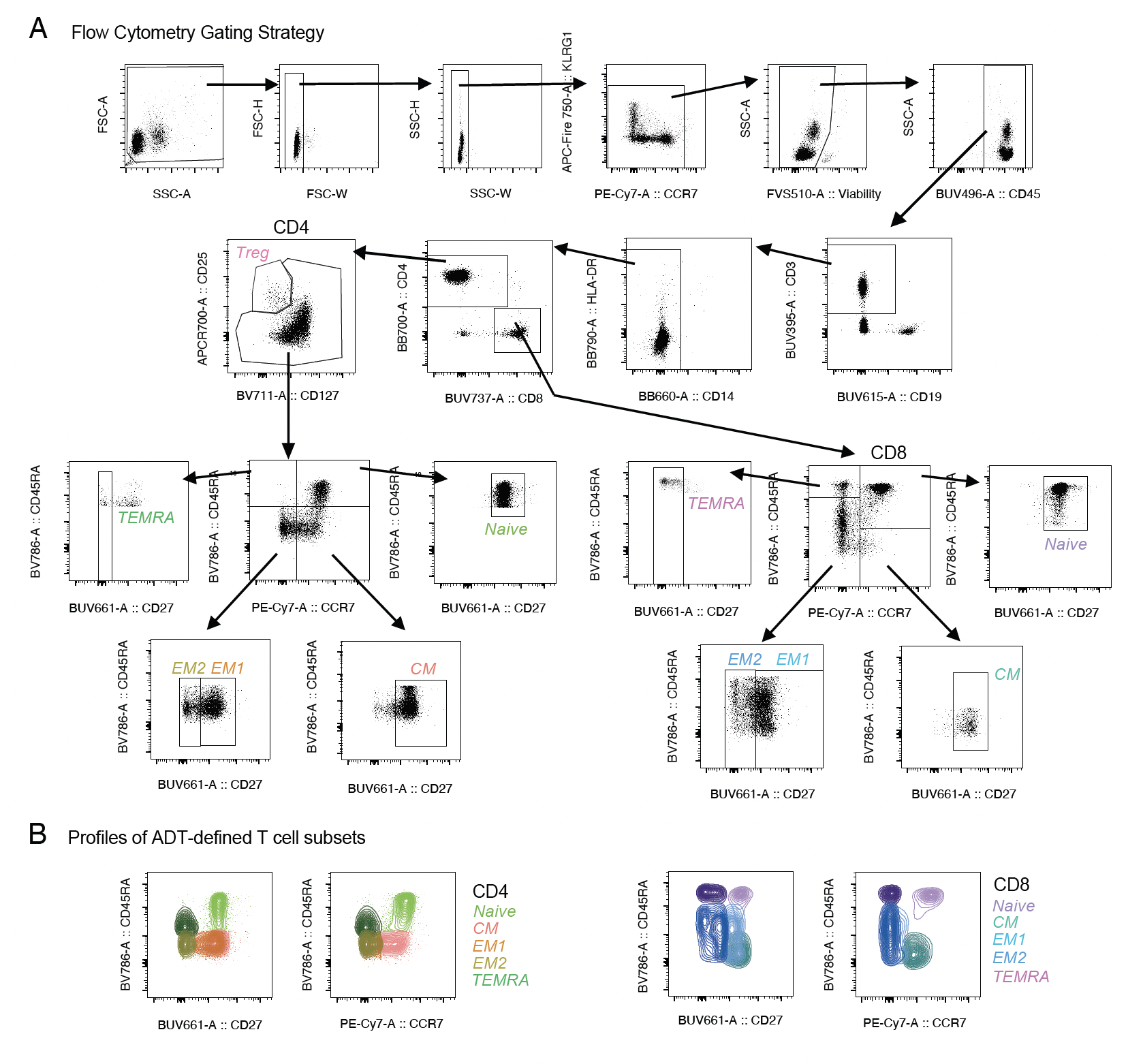

**Supplemental Figure 1**. **Flow cytometry gating of T cell subsets**. **(A)** Flow cytometry gating strategy for identifying T cell subsets. **(B)** Example contour plots of final T cell subsets based on strategy outlined in (A).

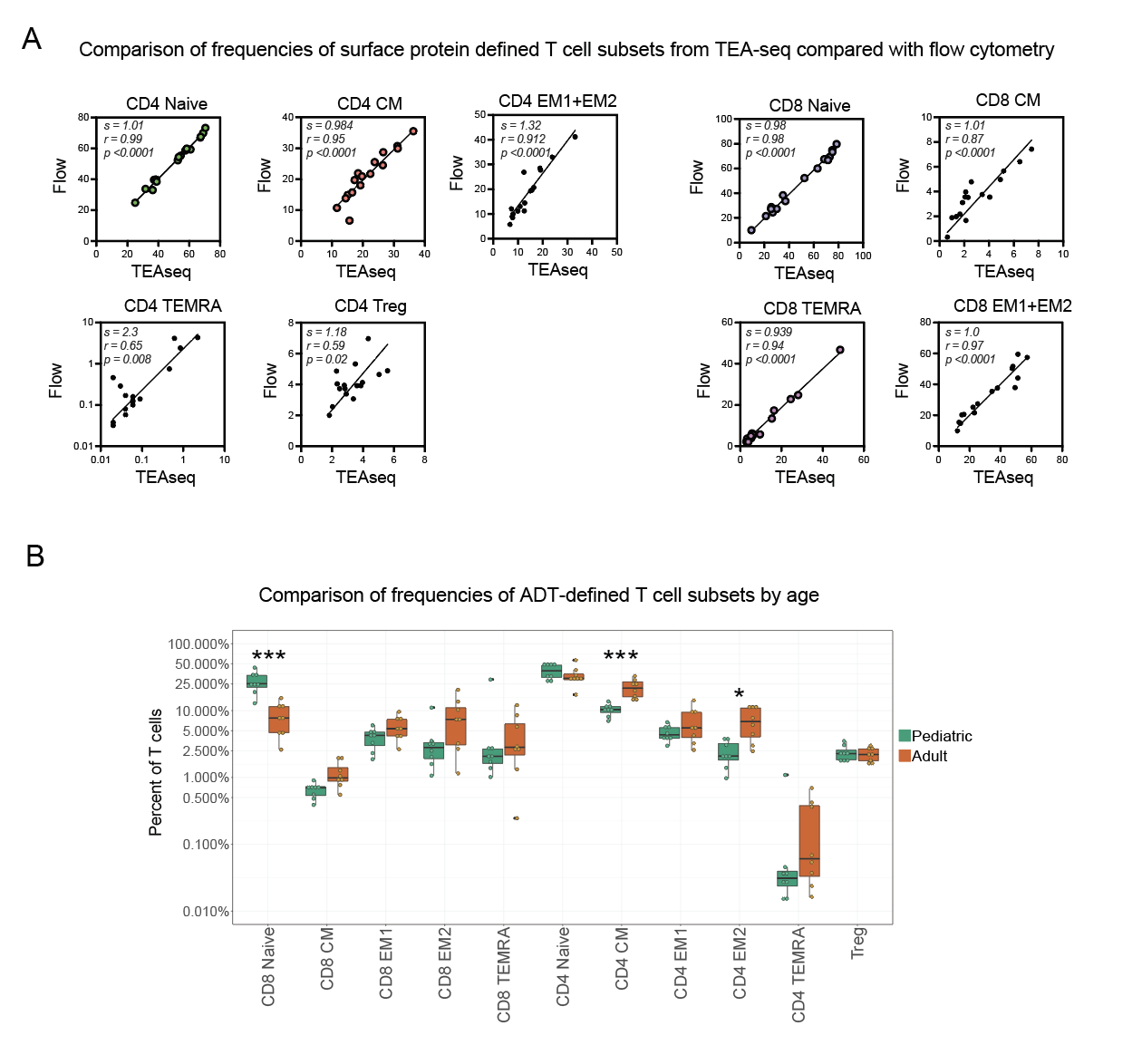

**Supplemental Figure 2**. **Comparison of ADT-defined T cell subsets in flow cytometry and age**. **(A)** Comparison of cell type proportions within total T cells identified by either TEA-seq or flow cytometry on the same donor samples. **(B)** Frequencies of ADT-defined T cell subsets within total T cell compartment by age group from TEA-seq. Mann-Whitney test with FDR multiple comparison analysis * q-value < 0.05, *** q < 0.001

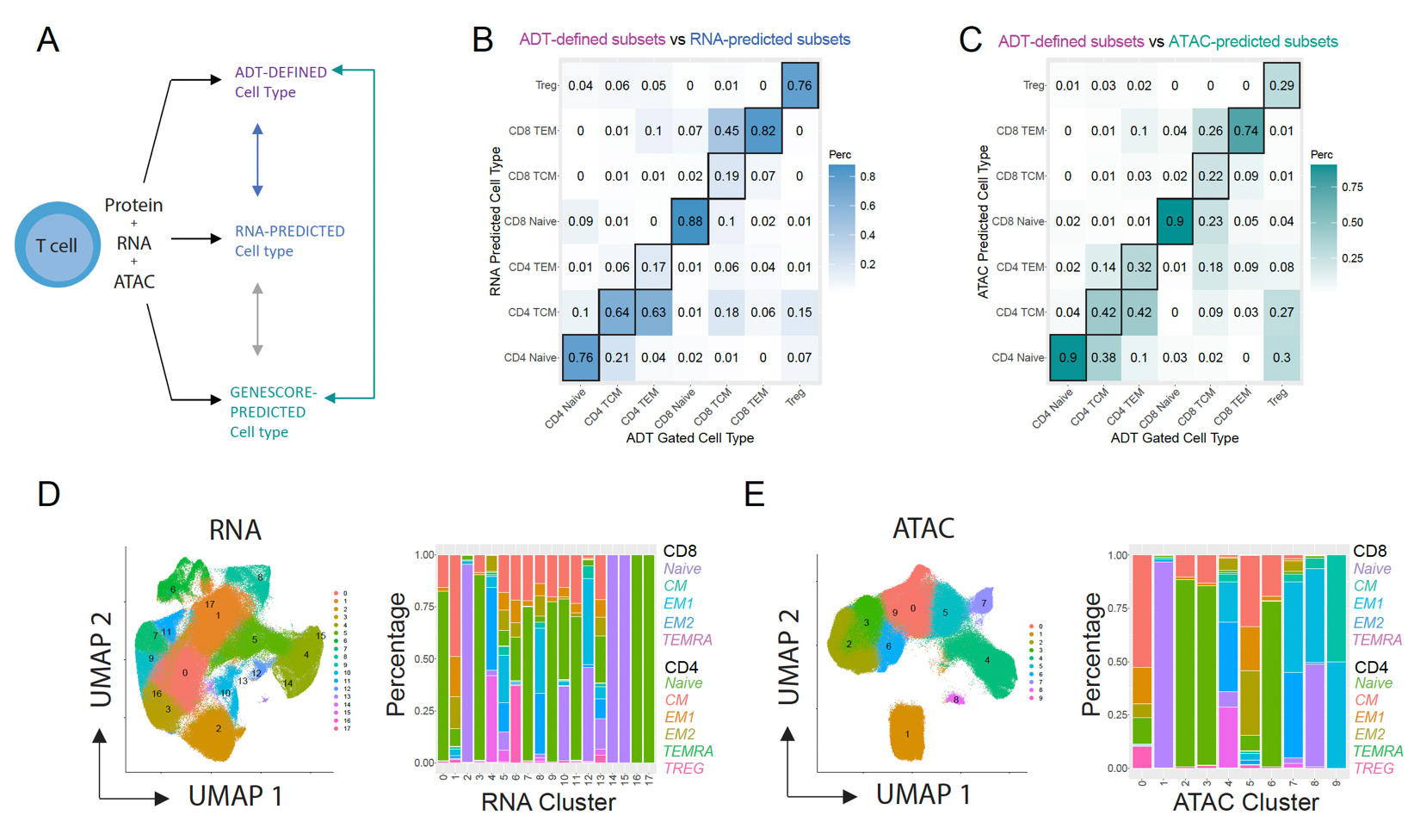

**Supplemental Figure 3. Cross-modal analysis of T cell subsets. (A)** Overview of single cell labeling methods used for each TEA-seq modality. **(B)** Confusion plot comparison of cell type labels between ADT-defined and RNA-based label transfer (Seurat). **(C)** Confusion plot comparison of cell type labels between ADT-defined and ATAC-based label transfer (ArchR). Values in **(B)** and **(C)** are the proportion of single cells defined by ADT that have the same cell type label in RNA or ATAC-based label transfer.

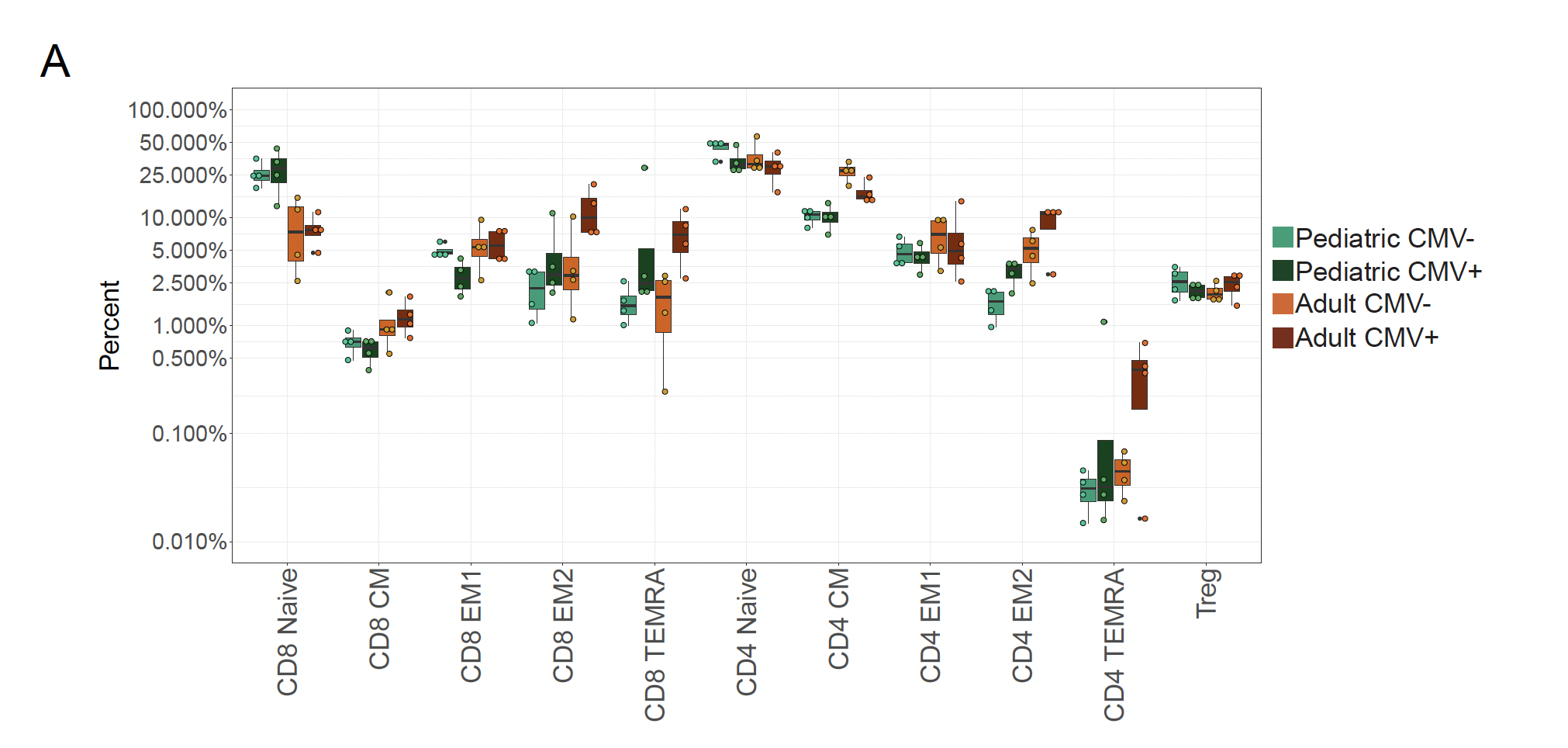

**Supplemental Figure 4. TEA-seq frequencies of T cell subsets by age and CMV infection status.** Naïve CD8 and CD4 central memory (CM) T cells show separation by age whereas TEMRA CD8 show separation by CMV infection status.

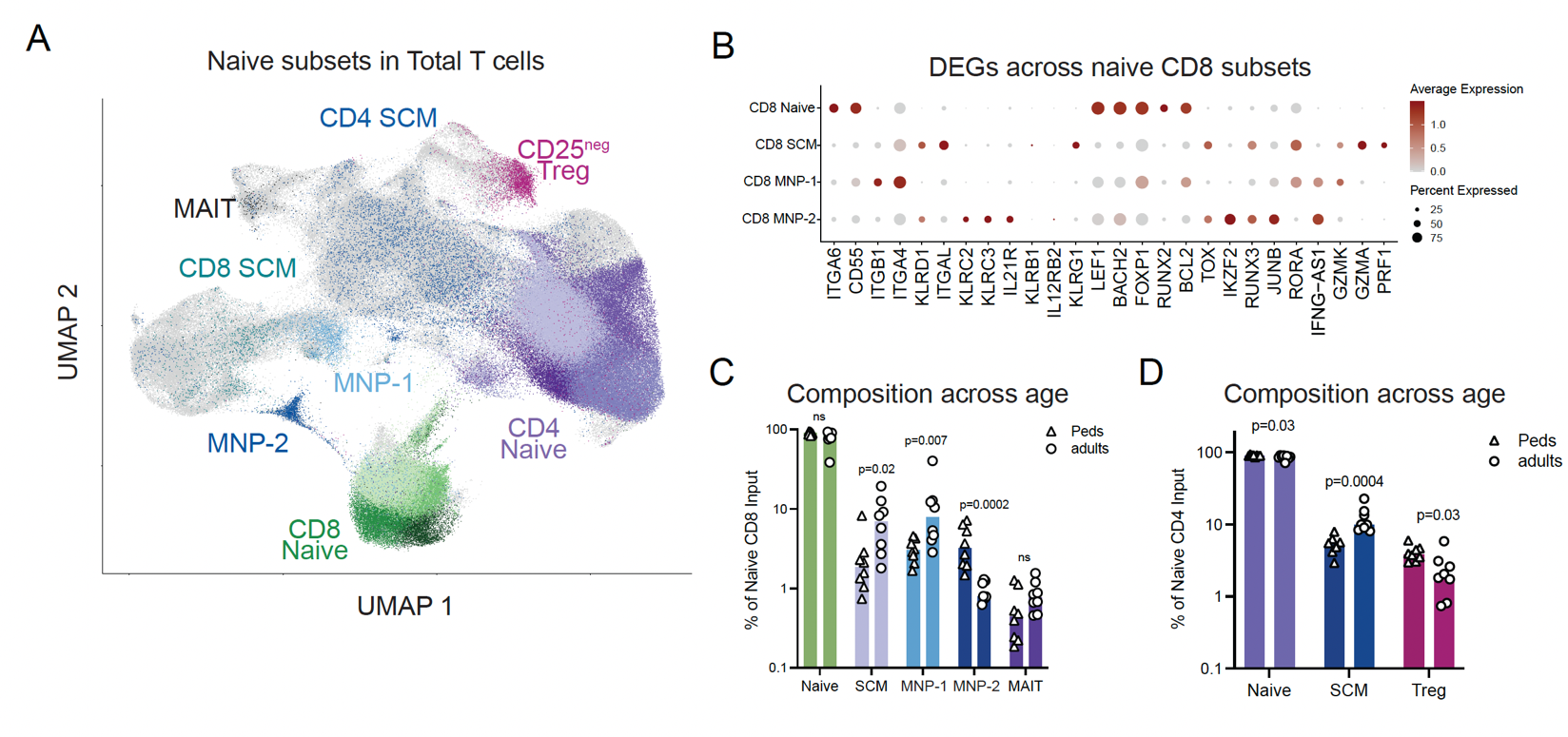

**Supplemental Figure 5. Select RNA profile of naïve CD8 T cell subsets.** Dot plot of select differentially expressed genes across naïve CD8 T cell subsets.

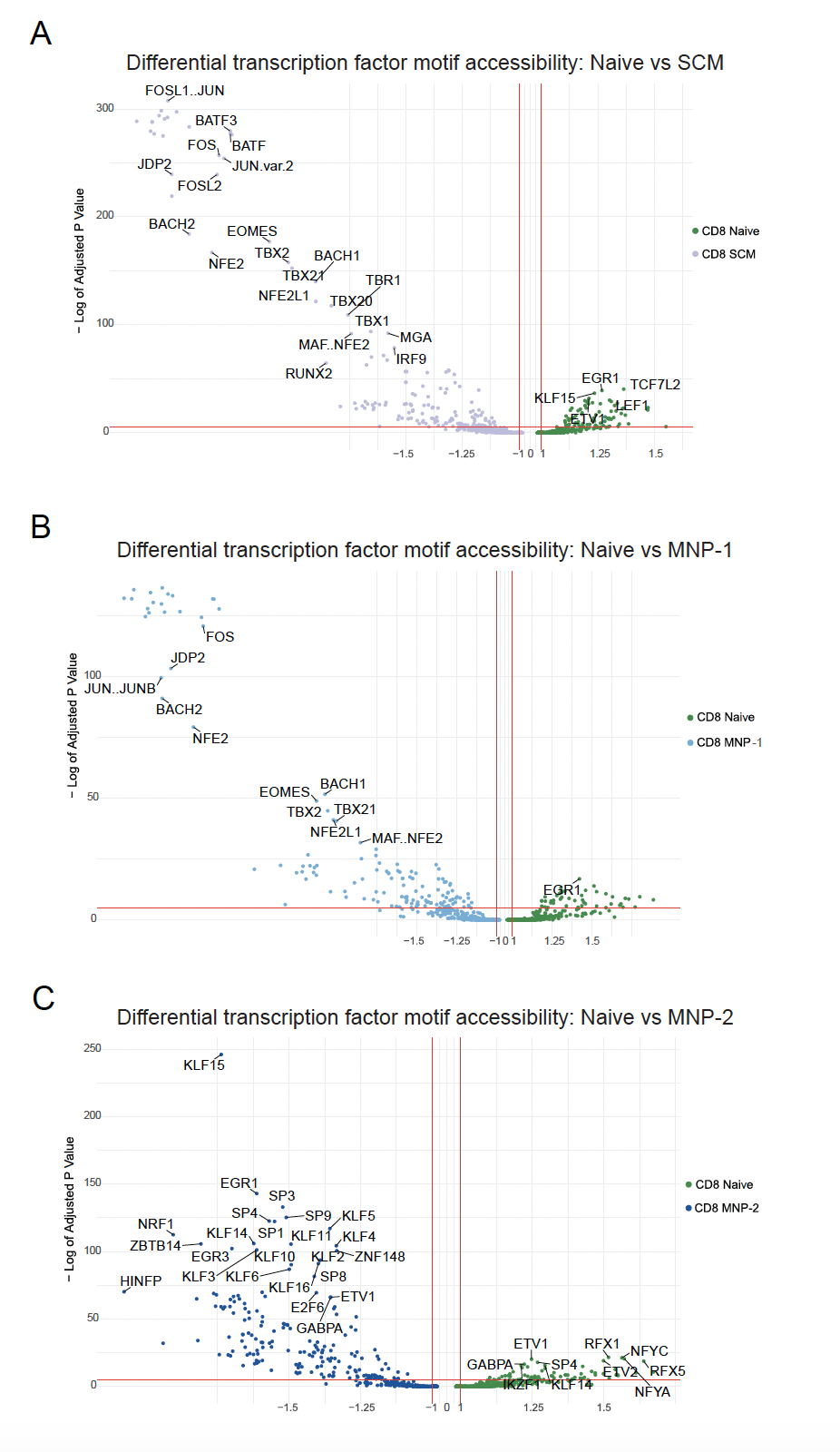

**Supplemental Figure 6**. **TF motif enrichment across naïve CD8 T cell subsets**. Transcription factor motif enrichment based on differentially accessible peaks between **(A)** true naïve vs SCM, **(B)** true naïve vs MNP-1 and **(C)** true naïve vs MNP-2 CD8 T cell subsets.

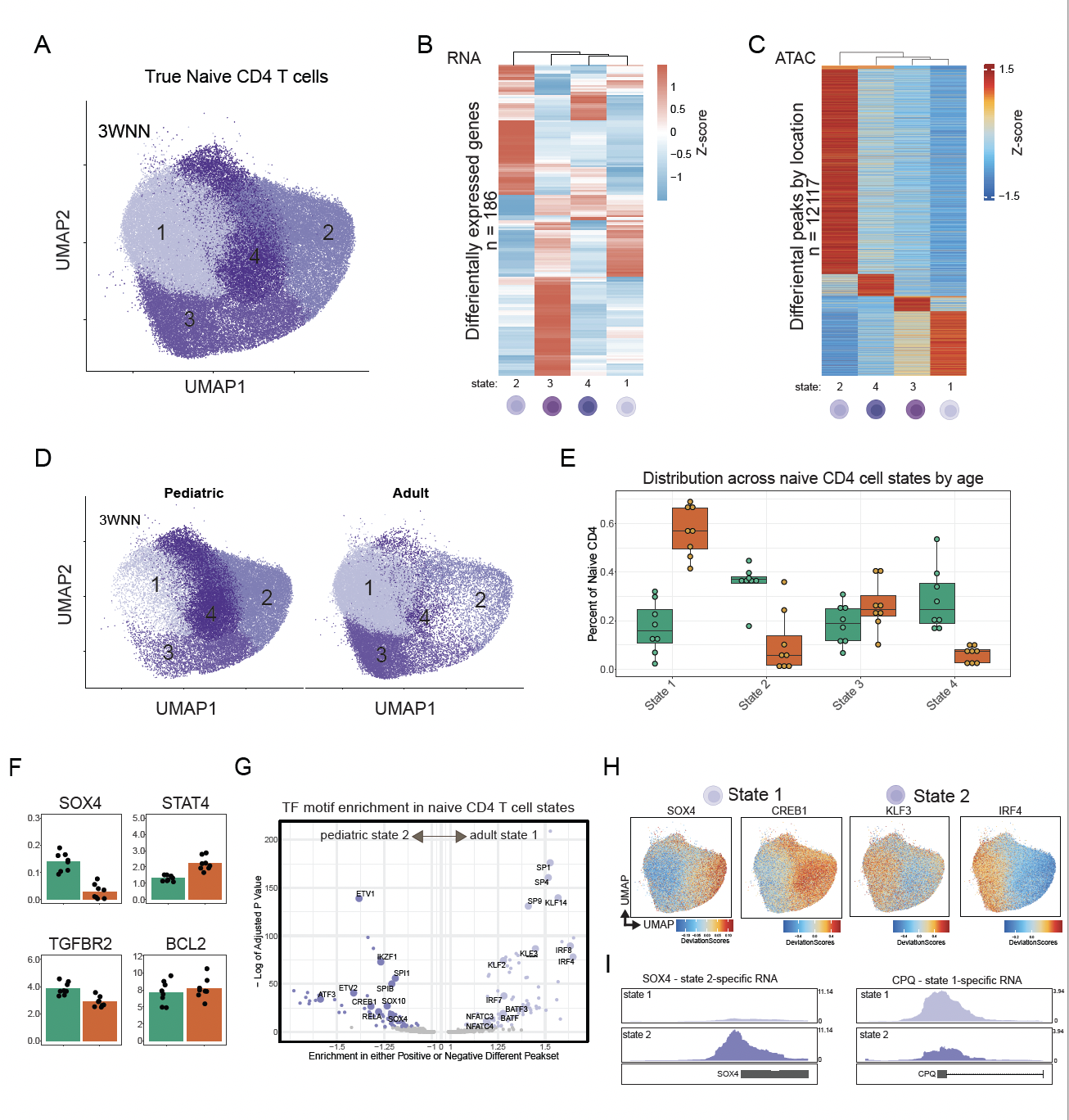

**Supplemental Figure 7**. Chromatin accessibility tracks of true naïve CD4 T cell state 1 and state 2-specific genes. Tracks are data from all true naïve CD4 T cells of children and adults combined.

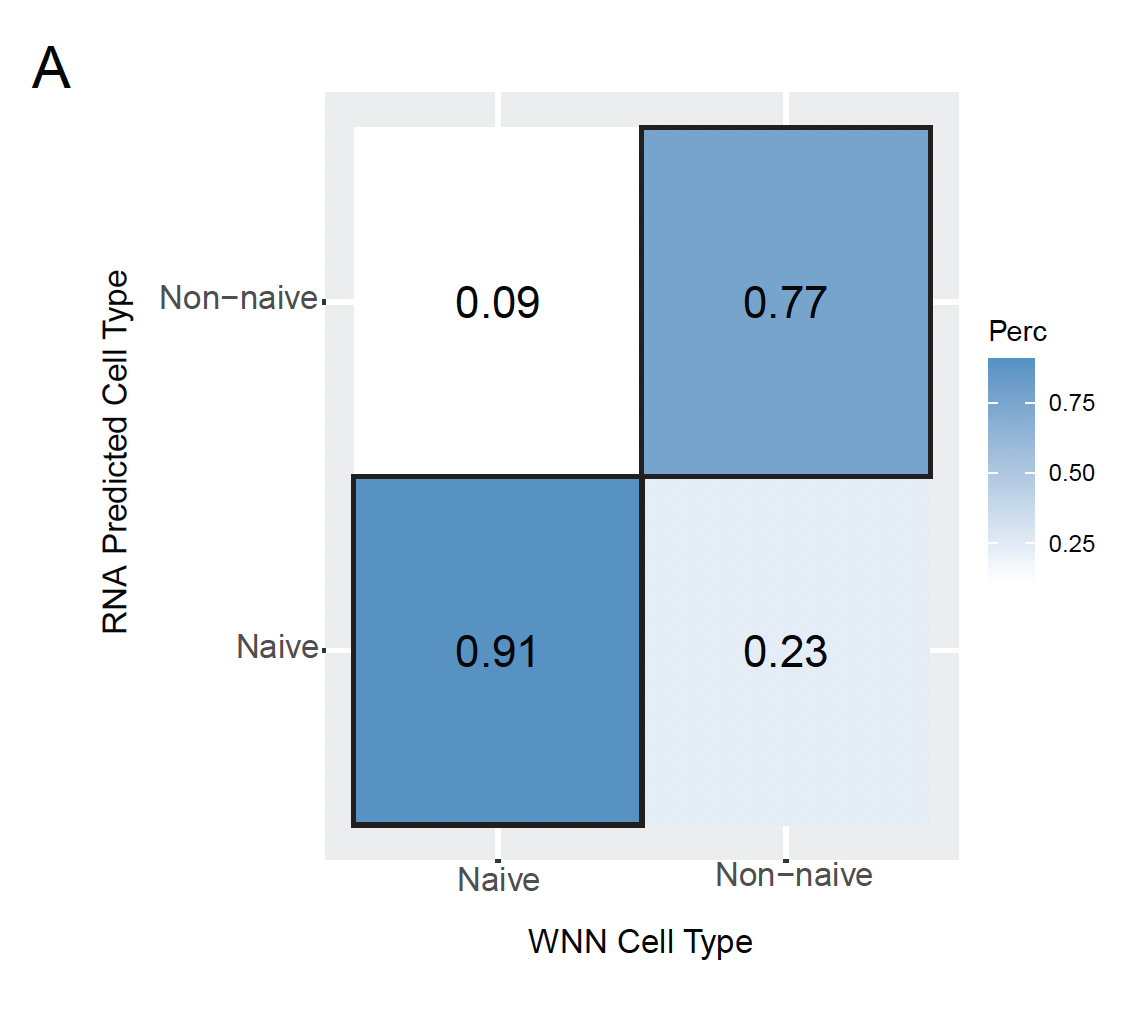

**Supplemental Figure 8**. **Confusion plot comparison** of WNN labels and RNA-based label transfer of ADT CD4+/CCR7+/CD45RA+/CD27+ cells.

**Supplemental Tables:**

**Supplemental Table 1: Cell Subset Markers for ADT-based Identification**

| **Cell Subset** | **ADT Markers** |
| --- | --- |
| CD4 Naïve | CD4+ / CD8- / CD197+ / CD27+ / CD45RA+ |
| CD4 CM | CD4+ / CD8- / CD197+ / CD27+ / CD45RA- |
| CD4 EM1 | CD4+ / CD8- / CD197- / CD27+ / CD45RA- |
| CD4 EM2 | CD4+ / CD8- / CD197- / CD27- / CD45RA- |
| CD4 TEMRA | CD4+ / CD8- / CD197- / CD27- / CD45RA+ |
| Treg | CD4+ / CD8- / CD127-/low / CD25+ |
| CD8 Naïve | CD4- / CD8+ / CD197+ / CD27+ / CD45RA+ |
| CD8 CM | CD4- / CD8+ / CD197+ / CD27+ / CD45RA- |
| CD8 EM1 | CD4- / CD8+ / CD197- / CD27+ / CD45RA- |
| CD8 EM2 | CD4- / CD8+ / CD197- / CD27- / CD45RA- |
| CD8 TEMRA | CD4- / CD8+ / CD197- / CD27- / CD45RA+ |

**Supplemental Table 2. Differentially expressed genes between Naïve CD8 T cell subsets.**

**Supplemental Table 3. Differentially expressed genes between true naïve CD4 T cell states.**

**Supplemental Table 4. Antibody List**

| **Reagent type (species) or resource** | **Designation** | **Clone (If applicable)** | **Source or reference** | **Identifiers** | **Additional Information** |
| --- | --- | --- | --- | --- | --- |
| **TEA-seq panel** | | | | | |
| Antibody | TotalSeq™-A0151 anti-human CD152 (CTLA-4) Antibody | BNI3 | BioLegend | Cat# 369619, RRID:AB_2734423 | (0.175 µg per million cells) |
| Antibody | TotalSeq™-A0071 anti-human CD194 (CCR4) Antibody | L291H4 | BioLegend | Cat# 359423, RRID:AB_2749979 | (0.175 µg per million cells) |
| Antibody | TotalSeq™-A0143 anti-human CD196 (CCR6) Antibody | G034E3 | BioLegend | Cat# 353437, RRID:AB_2750534 | (0.175 µg per million cells) |
| Antibody | TotalSeq™-A0189 anti-human CD244 (2B4) Antibody | C1.7 | BioLegend | Cat# 329527, RRID:AB_2750007 | (0.175 µg per million cells) |
| Antibody | TotalSeq™-A0396 anti-human CD26 Antibody | BA5b | BioLegend | Cat# 302720, RRID:AB_2734261 | (0.175 µg per million cells) |
| Antibody | TotalSeq™-A0102 anti-human CD294 (CRTH2) Antibody | BM16 | BioLegend | Cat# 350127, RRID:AB_2734360 | (0.2 µg per million cells) |
| Antibody | TotalSeq™-A0576 anti-human CD49d Antibody | 9F10 | BioLegend | Cat# 304337, RRID:AB_2783166 | (0.175 µg per million cells) |
| Antibody | TotalSeq™-A0171 anti-human/mouse/rat CD278 (ICOS) Antibody | C398.4A | BioLegend | Cat# 313555, RRID:AB_2800824 | (0.05 µg per million cells) |
| Antibody | TotalSeq™-A0161 anti-human CD11b Antibody | ICRF44 | BioLegend | Cat# 301353, RRID:AB_2734249 | (0.05 µg per million cells) |
| Antibody | TotalSeq™-A0053 anti-human CD11c Antibody | S-HCL-3 | BioLegend | Cat# 371519, RRID:AB_2749971 | (0.025 µg per million cells) |
| Antibody | TotalSeq™-A0390 anti-human CD127 (IL-7Rα) Antibody | A019D5 | BioLegend | Cat# 351352, RRID:AB_2734366 | (0.075 µg per million cells) |
| Antibody | TotalSeq™-A0083 anti-human CD16 Antibody | 3G8 | BioLegend | Cat# 302061, RRID:AB_2734255 | (0.05 µg per million cells) |
| Antibody | TotalSeq™-A0408 anti-human CD172a (SIRPα) Antibody | 15-414 | BioLegend | Cat# 372109, RRID:AB_2783285 | (0.25 µg per million cells) |
| Antibody | TotalSeq™-A0144 anti-human CD185 (CXCR5) Antibody | J252D4 | BioLegend | Cat# 356937, RRID:AB_2750356 | (0.125 µg per million cells) |
| Antibody | TotalSeq™-A0181 anti-human CD21 Antibody | Bu32 | BioLegend | Cat# 354915, RRID:AB_2750006 | (0.05 µg per million cells) |
| Antibody | TotalSeq™-A0085 anti-human CD25 Antibody | BC96 | BioLegend | Cat# 302643, RRID:AB_2734258 | (0.08 µg per million cells) |
| Antibody | TotalSeq™-A0154 anti-human CD27 Antibody | O323 | BioLegend | Cat# 302847, RRID:AB_2750000 | (0.05 µg per million cells) |
| Antibody | TotalSeq™-A0088 anti-human CD279 (PD-1) Antibody | EH12.2H7 | BioLegend | Cat# 329955, RRID:AB_2734322 | (0.2 µg per million cells) |
| Antibody | TotalSeq™-A0406 anti-human CD304 (Neuropilin-1) Antibody | 12C2 | BioLegend | Cat# 354525, RRID:AB_2783261 | (0.05 µg per million cells) |
| Antibody | TotalSeq™-A0410 anti-human CD38 Antibody | HB-7 | BioLegend | Cat# 356635, RRID:AB_2800967 | (0.05 µg per million cells) |
| Antibody | TotalSeq™-A0176 anti-human CD39 Antibody | A1 | BioLegend | Cat# 328233, RRID:AB_2750005 | (0.075 µg per million cells) |
| Antibody | TotalSeq™-A0072 anti-human CD4 Antibody | RPA-T4 | BioLegend | Cat# 300563, RRID:AB_2734247 | (0.1 µg per million cells) |
| Antibody | TotalSeq™-A0047 anti-human CD56 (NCAM) Antibody | 5.1H11 | BioLegend | Cat# 362557, RRID:AB_2749970 | (0.1 µg per million cells) |
| Antibody | TotalSeq™-A0394 anti-human CD71 Antibody | CY1G4 | BioLegend | Cat# 334123, RRID:AB_2800884 | (0.05 µg per million cells) |
| Antibody | TotalSeq™-A0080 anti-human CD8a Antibody | RPA-T8 | BioLegend | Cat# 301067, RRID:AB_2734248 | (0.2 µg per million cells) |
| Antibody | TotalSeq™-A0006 anti-human CD86 Antibody | IT2.2 | BioLegend | Cat# 305443, RRID:AB_2734273 | (0.05 µg per million cells) |
| Antibody | TotalSeq™-A0581 anti-human TCR Vα7.2 Antibody | 3C10 | BioLegend | Cat# 351733, RRID:AB_2783246 | (0.0625 µg per million cells) |
| Antibody | TotalSeq™-A0145 anti-human CD103 (Integrin αE) Antibody | Ber-ACT8 | BioLegend | Cat# 350231, RRID:AB_2749996 | (0.2 µg per million cells) |
| Antibody | TotalSeq™-A0168 anti-human CD57 Recombinant Antibody | QA17A04 | BioLegend | Cat# 393319, RRID:AB_2810588 | (0.2 µg per million cells) |
| Antibody | TotalSeq™-A0146 anti-human CD69 Antibody | FN50 | BioLegend | Cat# 310947, RRID:AB_2749997 | (0.2 µg per million cells) |
| Antibody | TotalSeq™-A0242 anti-human CD192 (CCR2) Antibody | K036C2 | BioLegend | Cat# 357229, RRID:AB_2750501 | (0.25 µg per million cells) |
| Antibody | TotalSeq™-A0063 anti-human CD45RA Antibody | HI100 | BioLegend | Cat# 304157, RRID:AB_2734267 | (0.25 µg per million cells) |
| Antibody | TotalSeq™-A0156 anti-human CD95 (Fas) Antibody | DX2 | BioLegend | Cat# 305649, RRID:AB_2750368 | (0.25 µg per million cells) |
| Antibody | TotalSeq™-A0159 anti-human HLA-DR Antibody | L243 | BioLegend | Cat# 307659, RRID:AB_2750001 | (0.05 µg per million cells) |
| Antibody | TotalSeq™-A0153 anti-human KLRG1 (MAFA) Antibody | SA231A2 | BioLegend | Cat# 367721, RRID:AB_2750373 | (0.25 µg per million cells) |
| Antibody | TotalSeq™-A0355 anti-human CD137 (4-1BB) Antibody | 4B4-1 | BioLegend | Cat# 309835, RRID:AB_2783173 | (0.25 µg per million cells) |
| Antibody | TotalSeq™-A0149 anti-human CD161 Antibody | HP-3G10 | BioLegend | Cat# 339945, RRID:AB_2749998 | (0.1 µg per million cells) |
| Antibody | TotalSeq™-A0140 anti-human CD183 (CXCR3) Antibody | G025H7 | BioLegend | Cat# 353745, RRID:AB_2749993 | (0.25 µg per million cells) |
| Antibody | TotalSeq™-A0896 anti-human CD85j (ILT2) Antibody | GHI/75 | BioLegend | Cat# 333723, RRID:AB_2814225 | (0.1 µg per million cells) |
| Antibody | TotalSeq™-A0179 anti-human CX3CR1 Antibody | K0124E1 | BioLegend | Cat# 355709, RRID:AB_2832698 | (0.1 µg per million cells) |
| Antibody | TotalSeq™-A0169 anti-human CD366 (Tim-3) Antibody | F38-2E2 | BioLegend | Cat# 345047, RRID:AB_2800924 | (0.2 µg per million cells) |
| Antibody | TotalSeq™-A0005 anti-human CD80 Antibody | 2D10 | BioLegend | Cat# 305239, RRID:AB_2749958 | (0.25 µg per million cells) |
| Antibody | TotalSeq™-A0148 anti-human CD197 (CCR7) Antibody | G043H7 | BioLegend | Cat# 353247, RRID:AB_2750357 | (0.5 µg per million cells) |
| Antibody | TotalSeq™-A0386 anti-human CD28 Antibody | CD28.2 | BioLegend | Cat# 302955, RRID:AB_2783159 | (0.5 µg per million cells) |
| Antibody | TotalSeq™-A0031 anti-human CD40 Antibody | 5C3 | BioLegend | Cat# 334346, RRID:AB_2749968 | (0.375 µg per million cells) |
| Antibody | TotalSeq™-A0087 anti-human CD45RO Antibody | UCHL1 | BioLegend | Cat# 304255, RRID:AB_2734268 | (0.5 µg per million cells) |
| Antibody | TotalSeq™-A0224 anti-human TCR α/β Antibody | IP26 | BioLegend | Cat# 306737, RRID:AB_2783167 | (0.375 µg per million cells) |
| Antibody | TotalSeq™-A0139 anti-human TCR γ/δ Antibody | B1 | BioLegend | Cat# 331229, RRID:AB_2734325 | (0.25 µg per million cells) |
| Antibody | TotalSeq™-A0089 anti-human TIGIT (VSTM3) Antibody | A15153G | BioLegend | Cat# 372725, RRID:AB_2734426 | (0.5 µg per million cells) |
| Antibody | TotalSeq™-A0158 anti-human CD134 (OX40) Antibody | Ber-ACT35 | BioLegend | Cat# 350033, RRID:AB_2783245 | (0.5 µg per million cells) |
| Antibody | TotalSeq™-A0032 anti-human CD154 Antibody | 24-31 | BioLegend | Cat# 310843, RRID:AB_2734283 | (0.5 µg per million cells) |
| Antibody | TotalSeq™-A0584 anti-human TCR Vα24-Jα18 (iNKT cell) Antibody | 6B11 | BioLegend | Cat# 342923, RRID:AB_2783227 | (0.5 µg per million cells) |
| Antibody | TotalSeq™-A0180 anti-human CD24 Antibody | ML5 | BioLegend | Cat# 311137, RRID:AB_2750374 | (0.5 µg per million cells) |
| Antibody | TotalSeq™-A0830 anti-human CD319 (CRACC) Antibody | 162.1 | BioLegend | Cat# 331821, RRID:AB_2800872 | (0.5 µg per million cells) |
| Antibody | TotalSeq™-A0090 Mouse IgG1, κ isotype Ctrl Antibody | MOPC-21 | BioLegend | Cat# 400199, RRID:AB_2868412 | (0.5 µg per million cells) |
| **Flow phenotyping panel** | | | | | |
| Antibody | Mouse anti-human CD3/BUV395 | UCHT1 | BD Bioscience | Cat# 563546, RRID:AB_2744387 |  |
| Antibody | Mouse anti-human CD45/BUV496 | HI30 | BD Bioscience | Cat# 624283 |  |
| Antibody | Mouse anti-human CD8/BUV737 | RPA-T8 | BD Bioscience | Cat# 624286 |  |
| Antibody | Mouse anti-human CD127/BV711 | A019D5 | BioLegend | Cat# 351328, RRID:AB_2562908 |  |
| Antibody | Mouse anti-human CD197/PE-Cy7 | G043H7 | BioLegend | Cat# 353226, RRID:AB_11126145 |  |
| Antibody | Mouse anti-human CD14/BB660 | MφP9 | BD Bioscience | Cat# 624295 |  |
| Antibody | Mouse anti-human CD56/BUV563 | NCAM16.2 | BD Bioscience | Cat# 612928 |  |
| Antibody | Mouse anti-human CD19/BUV615 | HIB19 | BD Bioscience | Cat# 624297 |  |
| Antibody | Mouse anti-human CD27/BUV661 | L128 | BD Bioscience | Cat# 624285 |  |
| Antibody | Mouse anti-human CD39/BUV805 | Tu66 | BD Bioscience | Cat# 624287 |  |
| Antibody | Mouse anti-human CD103/BV421 | Ber-ACT8 | BioLegend | Cat# 350214, RRID:AB_2563514 |  |
| Antibody | Mouse anti-human abTCR/BV480 | IP26 | BD Bioscience | Cat# 624278 |  |
| Antibody | Mouse anti-human CD223/BV605 | 11C3C65 | BioLegend | Cat# 369324, RRID:AB_2721541 |  |
| Antibody | Mouse anti-human CD95/BV650 | DX2 | BioLegend | Cat# 305642, RRID:AB_2632622 |  |
| Antibody | Mouse anti-human CD278/BV750 | DX29 | BD Bioscience | Cat# 624380 |  |
| Antibody | Mouse anti-human CD45RA/BV786 | HI100 | BioLegend | Cat# 304140, RRID:AB_2563816 |  |
| Antibody | Mouse anti-human CD185/BB515 | RF8B2 | BD Bioscience | Cat# 564624, RRID:AB_2738871 |  |
| Antibody | Mouse anti-human CD4/BB700 | SK3 | BD Bioscience | Cat# 566392, RRID:AB_2744421 |  |
| Antibody | Mouse anti-human HLA-DR/BB790 | G46-6 | BD Bioscience | Cat# 624296 |  |
| Antibody | Mouse anti-human CD279/PE | EH12.2H7 | BioLegend | Cat# 329906, RRID:AB_940483 |  |
| Antibody | Mouse anti-human TIGIT/PE-Dazzle594 | A15153G | BioLegend | Cat# 372716, RRID:AB_2632931 |  |
| Antibody | Mouse anti-human CD38/PE-Cy5 | HIT2 | BD Bioscience | Cat# 555461, RRID:AB_395854 |  |
| Antibody | Mouse anti-human CD69/APC | FN50 | BioLegend | Cat# 310910, RRID:AB_314845 |  |
| Antibody | Mouse anti-human CD25/APC-R700 | 2A3 | BD Bioscience | Cat# 565106, RRID:AB_2744339 |  |
| Antibody | Mouse anti-human KLRG1/APC-Fire750 | SA231A2 | BioLegend | Cat# 367718, RRID:AB_2687392 |  |
| **Flow sorting panel (Naïve CD4 T cells)** | | | | | |
| Antibody | Brilliant Violet 421™ anti-human CD95 (Fas) Antibody | DX2 | BioLegend | Cat# 305624, RRID:AB_2561830 |  |
| Antibody | FITC anti-human CD3 Antibody | UCHT1 | BioLegend | Cat# 300406, RRID:AB_314060 |  |
| Antibody | PerCP/Cyanine5.5 anti-human CD27 Antibody | O323 | BioLegend | Cat# 302820, RRID:AB_2073318 |  |
| Antibody | PE anti-human CD197 (CCR7) Antibody | G043H7 | BioLegend | Cat# 353204, 353204 |  |
| Antibody | PE-Cy™7 Mouse Anti-Human CD4 Antibody | SK3 | BD Bioscience | Cat# 557852, RRID:AB_396897 |  |
| Antibody | APC anti-human CD45RA Antibody | HI100 | BioLegend | Cat# 304112, RRID:AB_314416 |  |
| Antibody | APC/Cyanine7 anti-human CD8a Antibody | RPA-T8 | BioLegend | Cat# 301016, RRID:AB_314134 |  |
| **Flow sorting panel (Total T cells for TEA-seq)** | | | | | |
| Antibody | PE anti-human CD3 Antibody | UCHT1 | BioLegend | Cat# 300441, RRID:AB_2562047 |  |
| Antibody | FITC anti-human CD45 Antibody | HI30 | BioLegend | Cat# 304038, RRID:AB_2562050 |  |
